# Compound Delivery of eVLPs Enhances Prime Editing for Targeted Genome Engineering and High-Throughput Screening

**DOI:** 10.1101/2025.08.11.669692

**Authors:** Jethro Langley, Lou Baudrier, Jada Curry, Kiran Narta, Hayley Todesco, Kyle Potts, Sorana Morrissy, Douglas J. Mahoney, Pierre Billon

## Abstract

Engineered virus-like particles (eVLPs) enable transgene-free ribonucleoprotein delivery for genome editing applications, yet optimized delivery strategies for high-throughput applications remain unexplored. Prime editing enables precise genomic modifications but suffers from limited efficiency that constrains its widespread adoption. Here, we present PRIME-VLP (Progressive Repeated Infections for Maximized Editing via Virus-Like Particles), a delivery strategy that enhances prime editing efficiency for both targeted genome engineering and high-throughput prime editing screening. PRIME-VLP leverages the temporal dynamics of eVLP-mediated editing through multiple sequential transductions with sub-saturating eVLP doses delivered at optimal intervals. This approach achieves 1.5 to 2.8-fold improvements in editing efficiency across diverse genomic targets and cell types. PRIME-VLP maintains high specificity without increasing off-target effects, compromising cellular viability or causing transcriptional perturbations. By decoupling pegRNA and editor delivery through pegRNA-free eVLPs, PRIME-VLP enables pooled prime editing screens, circumventing transgene silencing limitations of conventional lentiviral-based screens. Using a 6,000-pegRNA library targeting *TP53*, PRIME-VLP achieved 2.8-fold higher editing efficiency and improved reproducibility compared to conventional lentiviral delivery. An eVLP-based screen identified functional *TP53* loss-of-function variants that confer resistance to MDM2 inhibition by Nutlin-3. This work expands the versatility of eVLPs beyond their current *in vivo* therapeutic applications, demonstrating their promise for high-throughput functional genomics by overcoming the delivery limitations of lentiviral systems.

## INTRODUCTION

The exponential growth in genomic sequencing has enabled the identification of genetic variants across diverse populations and disease contexts^1–4^, yet the majority remain variants of uncertain significance due to the lack of functional interpretation, which has restricted their clinical application^5,6^. Early functional screening efforts relied on ectopic cDNA overexpression systems to assess variant function but were limited by non-physiological expression patterns and cellular contexts. The development of high-throughput screening platforms using pooled short-hairpin RNA libraries enabled specific perturbation of gene transcripts using lentivirally-delivered libraries^7,8^. In the past decade, CRISPR-based high-throughput screens have emerged as a powerful approach for elucidating gene function at scale within their native genomic context^9,10^. The proven success of lentiviral delivery in cDNA and RNAi screens naturally led to its adoption for CRISPR screening applications, where lentiviruses became the standard method for delivering both genome editors and guide RNA (gRNA) libraries. Lentiviruses enable controlled transgene dosing and provide stable transgene integration that is maintained through cell divisions, making them well-suited for large-scale pooled screening applications.

However, functionally characterizing individual genetic variants requires transitioning from gene-level perturbations to achieve single-nucleotide precision. Recent advances in precision genome editing technologies, such as base editing^11,12^ and prime editing^13^, have enabled systematic evaluation of genetic variants. Prime editing is particularly promising as it enables the precise installation of all types of base substitutions, insertions or deletions, without requiring double-strand breaks or donor templates^14,15^. This versatility offers a unique opportunity to functionally characterize variants that remain to be linked to human variation or diseases. Despite this versatility and potential for functional genomics, the widespread adoption of prime editing in precision genome editing and high-throughput applications has been constrained by low editing efficiency^13,14^. Recent optimizations and engineering have been developed to enhance prime editing efficiency, including optimizing pegRNA architecture^16–18^, introducing nicks in the non-edited strand^13^, transiently inhibiting repair pathways^19,20^, engineering enhanced prime editing components^20–23^, modulating cellular metabolism^24^ and implementing co-selection approaches^25,26^. Although these advances have enhanced prime editing performance, achieving reproducible high editing across thousands of targets in large-scale functional screens remains challenging.

One of the primary reasons for this challenge is the difficulty of efficiently delivering large prime editing components to target cells in current screening methodologies. Given the low efficiency of prime editing, it is essential to maintain stable, high-level expression of the prime editor to accumulate edits over time, which can require extended editing periods up to a month^27^. Existing approaches rely predominantly on lentiviral integration^28–31^ of large editor constructs. However, lentiviral delivery suffers from three major limitations: (1) variable transgene expression due to semi-random integration sites^31,32^, (2) progressive silencing through promoter methylation and establishment of chromatin repression marks over time^33,34^, and (3) packaging limits that result in low functional titers^35–37^ or require the development of complex multi-vector strategies^13^. These limitations lead to inconsistent editing across cell populations and poor experimental reproducibility. Recent prime editing screens^26–31,38,39^ have highlighted these delivery challenges, necessitating extensive optimization strategies, including site-specific integration of editor^27^ and single-cell cloning to isolate high-expressing populations^26,30,31,38,39^, disrupting mismatch repair^27^, or the use of co-selection methods^26^. The lack of an effective delivery platform represents a critical bottleneck preventing the widespread adoption of prime editing for functional genomics.

These delivery limitations highlight the need for delivery approaches that bypass transgene integration entirely. Engineered virus-like particles (eVLPs) represent an emerging modular RNP delivery system with unique advantages for genome editing applications^40–46^. Recent studies have demonstrated that eVLPs can efficiently deliver Cas9^40,43–45^, base editors^40,46^ and prime editors^41,46^ to different cell types and animal tissues^40,41,46^. Unlike conventional lentiviral vectors, eVLPs directly package and deliver functional ribonucleoprotein (RNP) complexes by fusing editing machinery to viral structural proteins for efficient packaging during particle assembly. This approach circumvents the limitations of transgene integration and expression by delivering pre-assembled RNP complexes that begin functioning immediately upon cellular entry, without requiring transcription or translation. eVLPs offer several advantages that can benefit high-throughput applications, which include their self-assembly mechanism that enables scalable production in standard laboratory settings, capacity to package large protein complexes, compatibility with various viral architectures, such as MMLV^40,41^ or HIV^43,44,46^ or engineered viral components, such as miniaturized EDV (mini-EDVs)^47^ or eVLPv5^42^, and flexible tropism that allows retargeting to specific cell types^40,43–45,48^. While eVLPs hold considerable therapeutic potential, their application in high-throughput CRISPR-based screens has remained unexplored. Key questions concerning their efficiency for large-scale editor delivery, compatibility with established screening platforms, and potential unintended effects on cellular physiology remain unresolved.

To overcome these delivery constraints, we developed a novel eVLP-based delivery method that leverages the temporal dynamics of RNP-based editing. We first performed analysis of eVLP-mediated editing kinetics and dose-response relationships, providing the mechanistic rationale for optimally timed sequential deliveries that maintain steady-state editor levels while avoiding saturation effects. This led to the development of PRIME-VLP, a compound delivery strategy that uses multiple temporally spaced, sub-saturating eVLP doses to overcome saturation limitations to significantly enhance prime editing efficiency. Our studies show that PRIME-VLP consistently achieves 1.5-2.8-fold improvements in prime editing efficiency across diverse genomic targets and cell types without increasing off-target effects, compromising cellular viability or disrupting gene expression profiles. To enable pooled screening applications, we further developed a decoupling strategy that separates editor and pegRNA delivery, circumventing the transgene silencing and heterogeneous expression limitations that limit current screening approaches. We demonstrate the practical utility of this approach through a 6,000-pegRNA library screen targeting *TP53*, achieving 2.8-fold superior performance compared to conventional lentiviral delivery and identification of *TP53* loss-of-function variants that confer resistance to Nutlin-3. This work expands the versatility of eVLPs beyond their current *in vivo* therapies, demonstrating their promise for functional genomics by overcoming the delivery limitations of lentiviral systems.

## RESULTS

### Development and characterization of eVLPs for precision genome editing

To evaluate the potential of eVLPs as a delivery platform for precision genome editing, we first adapted a previously described Moloney Murine Leukemia Virus (MMLV) Gag-fusion approach using an adenine base editor (ABE8e)^40^ to package additional CRISPR-based genome editors, including a cytosine base editor (tadCBEd) and a prime editor (PE2max) (**Figure 1A-1C, left**). We produced these eVLPs using previously published protocols^40,41^, yielding a 100 μL preparation from the standard production scale. Tunable resistive pulse sensing (TRPS) quantification of eVLP preparations revealed consistent particle concentrations across editor types (average: 4 x 10^12^ particles/mL, range: 1.75-7.00 x 10^12^ particles/mL, n = 9) with PE2max-eVLPs showing particularly tight reproducibility (range: 2.92-3.41 x 10^12^ particles/mL). Modal particle diameters consistently centered between 100-150 nm across all editor types, consistent with previous observations of similar particle types^40^ (**Figure S1A and S1B**). This production consistency highlights the reproducibility and robustness of established eVLP production protocols (see **Materials and Methods**). Having established consistent eVLP production and packaging, we next sought to validate their editing performance.

**Figure 1.**
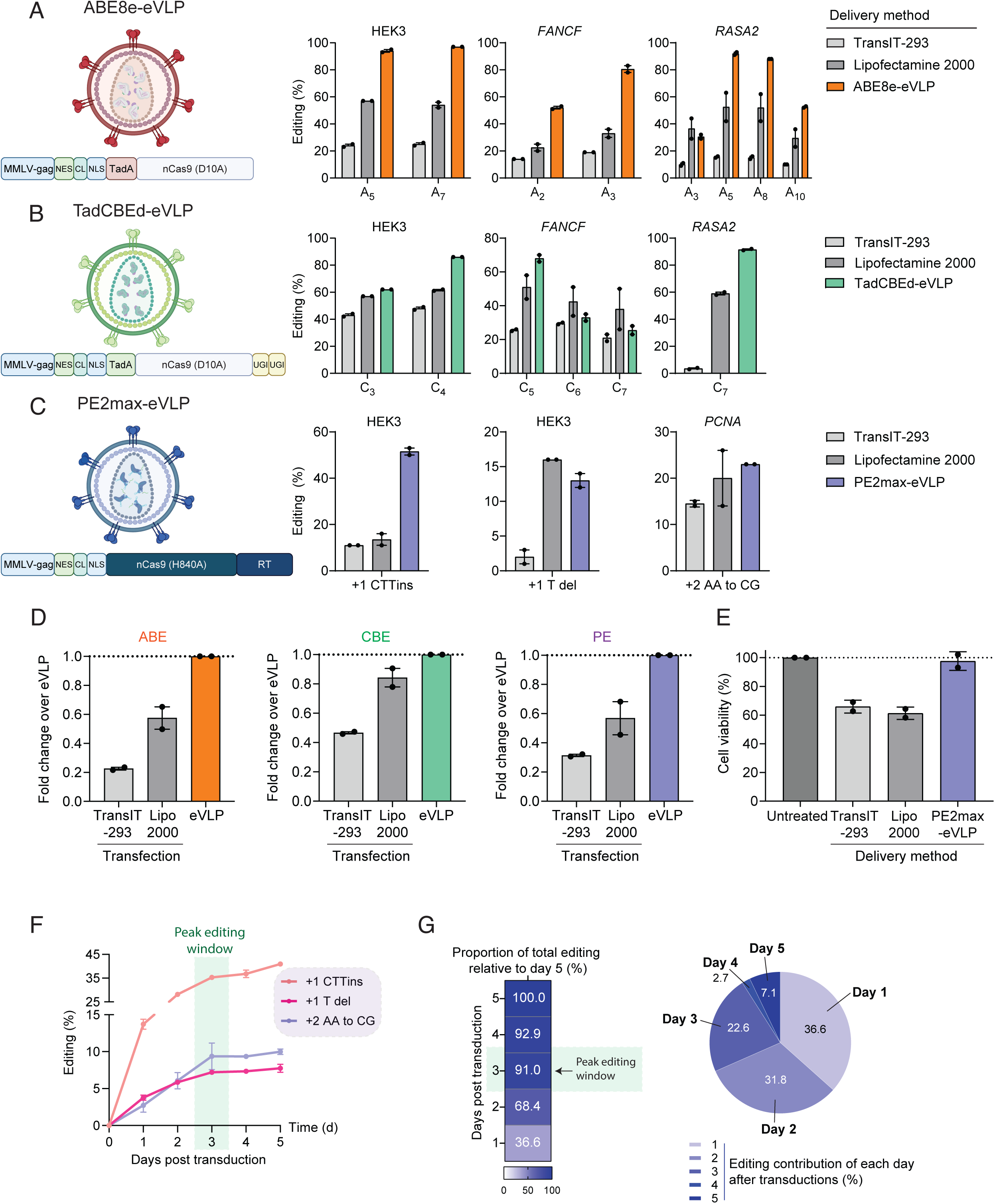
Development and characterization of eVLPs for precision genome editing. **(A-C)** Schematic representations and benchmarking of eVLPs packaging different genome editors. Left: eVLP architecture with MMLV fusion proteins packaging adenine base editors (ABE8e, orange), cytosine base editors (TadCBEd, green), or prime editors (PE2max, purple). Right: Editing efficiency comparison between eVLP delivery and standard transfection methods (*Trans*IT-293 and Lipofectamine 2000) at endogenous genomic loci in HEK293T cells. Data represent mean ± s.e.m from n = 2 independent experiments. NES: Nuclear Export Sequence, CL: Cleavable Linker, NLS: Nuclear Localization Sequence, UGI: Uracil Glycosylase Inhibitor, RT: Reverse Transcriptase. **(D)** Fold-change analysis of eVLP delivery efficiency relative to transfection methods (normalized to 1.0, dotted line) across adenine base editing (ABE), cytosine base editing (CBE), and prime editing (PE) platforms. Data represent mean ± s.e.m. (n=2). **(E)** Cell viability assessment 72 hours post-treatment comparing untreated controls with each delivery method. Data represent mean ± s.e.m. (n=2). **(F)** Kinetics analysis of prime editing efficiency following PE2max-eVLP transduction. Three independent genomic loci (HEK3 +1 CTTins, HEK3 +1 T del, *PCNA* +2 AA to CG). Green shading highlights the peak editing window (day 3). **(G)** Temporal analysis of editing kinetics. Left: Heat map displaying cumulative proportion of total editing achieved by each day relative to final day 5 values. Right: pie chart showing daily contribution of each time point to the total editing observed by day 5, demonstrating that 91% editing occurs within the first 3 days.

We benchmarked eVLP editing performance against conventional plasmid transfection using TransIT-293 and Lipofectamine 2000, following manufacturer and published protocols^49,50^. We used a full 100 μL eVLP preparation (corresponding to 4 x 10^11^ particles) to ensure maximal editing for accurate benchmarking against transfection methods. While studies typically employ 0.5-50 μL doses for precision genome editing^40,41^, we used the maximum available dose to establish optimal editing conditions before investigating dose optimization strategies. In HEK293T cells, eVLPs consistently outperformed both transfection methods across multiple genomic loci and edit types (**Figures 1A-1D and S1C**). While transfection achieved editing efficiencies of 12.7-31% across editors (ABE: 29.3%, CBE: 31.3%, PE: 12.7%), eVLPs delivered significantly higher efficiencies (ABE: 73.3%, CBE: 61%, PE: 29.2%), representing 2.0 to 2.5-fold improvements. Notably, eVLPs maintained high cell viability profiles, while transfection reagents reduced viability by approximately 40% at 72 hours post-transfection (**Figure 1E**). These results demonstrate that eVLPs can efficiently deliver highly functional ABE8e, tadCBEd, and PE2max editing machinery to target cells, which is consistent with independent work^40,41^.

To characterize eVLP-mediated editing kinetics and identify potential optimization opportunities, we conducted kinetic analysis of prime editing across three independent genomic loci. Cells were transduced with PE2max-eVLPs and harvested at daily intervals over five days, with editing frequencies quantified by next-generation sequencing (NGS). Notably, all three loci exhibited remarkably similar kinetics: rapid increase in editing efficiency over the first two days post-transduction, followed by a plateau phase reached by day 3 (**Figure 1F**). Quantitative analysis of the relative contribution of each day to total editing revealed that the vast majority of edits (91%, n =3) occurred within the first 3 days (**Figure 1G**), with individual loci reaching 86.3%, 93.1%, and 93.6% of the total editing observed on day 5 (**Figure S1D**). This rapid editing profile, which is consistent with the short half-life of RNP complexes in cells^51^, and DNA repair kinetics^52^, indicates that maximal editing occurs within a defined three-day window post-transduction. Given that prime editing efficiency can benefit from sustained editor activity through high expression of prime editors for prolonged periods^27,29,53,54^, our findings raise the possibility that sequential dosing, optimally timed to coincide with the decline of the initial editing activity, could further enhance overall editing efficiency.

### Compound delivery of eVLPs overcomes editing saturation and enhances prime editing efficiency

Building on our observation that eVLP-mediated editing reaches peak efficiency within three days before plateauing, we hypothesized that sequential eVLP transductions could maintain sustained editor activity and enhance overall editing efficiency. To optimize a dosing strategy for sequential PE2max-eVLP transductions, we first characterized the dose-response relationship between eVLP input and editing efficiency. To assess dose-dependent effects, we performed a titration across three targets using PE2max-eVLPs ranging from 5 μL to 100 μL (**Figure 2A**). Editing efficiency at 3 days post-transduction demonstrated non-linear dose-dependent responses at all three genomic loci (**Figure 2B and S2A-S2C**). Editing efficiency increased sharply from an average of 9.4% at the lowest dose to 20.9% at intermediate doses, but plateaued at 24.5%, despite further dose escalation to 100 μL. Critically, at 25 μL, editing reached 92.9% of the efficiency observed at 100 μL, indicating that sub-saturating doses achieve near-maximal editing (**Figure 2C**). This saturation effect was observed consistently across all targeted loci despite their different maximum editing efficiencies.

**Figure 2.**
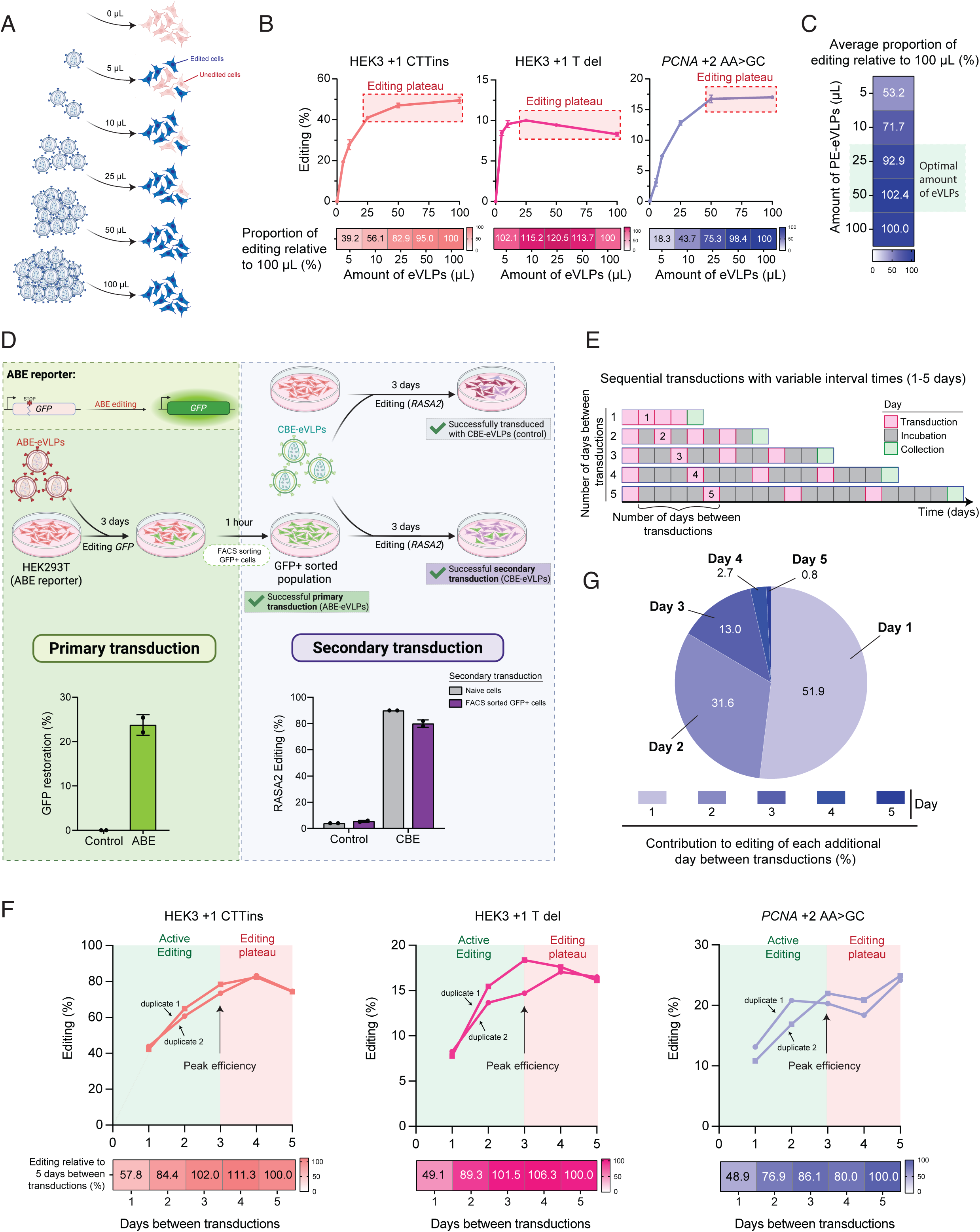
Sequential eVLP transductions overcome saturation and improve prime editing efficiency. **(A)** Experimental design for dose-response analysis using increasing eVLP volumes (5-100 μL). Schematic illustrates eVLP particles (blue circles) transducing cell populations, with blue cells representing edited cells and pink cells representing unedited cells. Increasing eVLP doses result in progressively higher proportions of edited cells. **(B)** Dose-response curves for PE2max-eVLPs across three genomic targets demonstrating saturation plateau at higher doses. Heat maps below show editing efficiency values for each dose tested. Red dashed boxes highlight the saturation plateau region. **(C)** Quantification of editing efficiency relative to the 100 μL dose (set to 100%) reveals that 25 μL achieves 92.9% of maximal editing across all targets, establishing the optimal sub-saturating dose for sequential transductions. **(D)** Sequential transduction validation experiment demonstrating cellular permissiveness to multiple eVLP deliveries. ABE-eVLPs restore GFP expression in reporter cells (primary transduction), followed by TadCBEd-eVLP targeting RASA2 in sorted GFP+ cells (secondary transduction). Bar graphs show successful primary transduction (23.8% GFP restoration, left) and comparable secondary editing efficiency in pre-transduced GFP+ cells versus naive control cells (right). Data represent mean ± s.e.m. from n = 2 independent experiments. **(E)** Experimental design for optimizing number of days between sequential transductions (1-5 days). Four sequential PE2max-eVLP transductions were performed with variable intervals (1-5 days) to determine optimal timing. Color coding indicates transduction events (purple), incubation periods (grey), and collection timepoints (green). **(F)** Prime editing efficiency optimization across different inter-transduction intervals for three independent genomic loci. Graphs show editing efficiency versus days between transductions, with arrows indicating peak efficiency for each replicate. Green shading highlights the active editing phase (1-3 days) while pink shading indicates the saturation phase (4-5 days) where additional time provides diminishing returns. **(G)** Pie chart displays the percentage contribution of each additional day between transductions to total editing improvement, demonstrating that 1-3 day intervals provide 51.9%, 31.6%, and 13.0% of maximal editing benefit, respectively, while longer intervals (days 4-5) show minimal additional benefit (2.7% and 0.8%).

Given that eVLP-mediated editing exhibits both defined temporal kinetics (**Figure 1F and 1G**) and dose-dependent saturation (**Figure 2A-2C**), we next investigated whether sequential transductions could overcome these constraints. We reasoned that multiple, temporally spaced, sub-saturating transductions could enhance editing efficiency by maintaining steady-state editor levels while avoiding saturation-induced plateaus. Since eVLPs do not contain integrative nucleic acids, they should bypass cellular innate immune mechanisms that typically prevent viral superinfection^55^. To validate this concept, we employed a two-step editing approach to sequentially edit two genomic loci (**Figure 2D**). First, HEK293T cells harbouring a GFP reporter with a premature stop codon were transduced with ABE-eVLPs containing a gRNA that restored GFP expression upon successful editing. This initial transduction resulted in 23.8% GFP restoration (**Figure 2D, left**), confirming successful transduction. GFP+ cells were subsequently sorted to isolate a population that had successfully undergone primary transduction. Immediately following sorting, cells were re-transduced with tadCBEd-eVLP carrying a gRNA targeting the endogenous *RASA2* gene. We used tadCBEd for the secondary transduction to avoid potential confounding effects from residual ABE activity from the initial transduction. Notably, *RASA2* editing efficiency was highly comparable between the secondary infection (80%) and primary transduction of naïve cells (90%) (**Figure 2D, right**), demonstrating that cells remain permissive to eVLP retransduction after initial editing. We confirmed these findings in primary human T cells by comparing single-dose to sequential dosing with tadCBEd-eVLPs. Sequential dosing improved editing efficiency, indicating that primary cells with intact innate immune responses remain permissive to eVLP-mediated editing (**Figure S2D**). These results establish the feasibility of sequential transductions with eVLPs without loss of cellular permissiveness.

Having confirmed that sequential transductions are feasible, we next optimized the timing intervals between these eVLP transductions to maximize cumulative editing efficiency. To determine optimal timing between sequential transductions, cells were subjected to four rounds of PE2max-eVLP transductions, varying the transduction interval from 1 to 5 days (**Figure 2E**). Editing efficiency increased progressively with longer intervals up to three days (**Figure 2F**, green shading), beyond which additional days provided little additional benefit (**Figure 2F**, pink shading). Specifically, intervals of 1-3 days between transductions contributed 51.9%, 31.6%, and 13% of the maximal editing, respectively (**Figure 2G**). In contrast, days 4 and 5 contributed only 2.7% and 0.8%, indicating a plateau in cumulative editing frequency beyond the three-day interval (**Figure 2G**). These results confirmed that peak efficiency occurs with three-day intervals, providing sufficient time for each round of editing to complete while maintaining optimal efficiency. This timing precisely aligns with our kinetics data showing that editing reaches maximum levels within 3 days (**Figure 1F and 1G**), providing a mechanistic foundation for our compound delivery approach.

Together, these studies identified key parameters for enhanced eVLP-mediated editing: sub-saturating doses (25 μL) delivered at three-day intervals to maintain sustained editor activity while avoiding saturation effects.

### PRIME-VLP improves editing efficiency through compound eVLP transductions

To evaluate PRIME-VLP performance, we compared prime editing efficiencies across six endogenous genomic targets representing diverse edit types. Cells received either a single saturating dose of eVLPs (one transduction of 100 μL, ∼4 x 10^11^ particles) or the same total number of particles delivered via the PRIME-VLP strategy (four sequential transductions of 25 μL each). To eliminate technical variability between conditions, eVLPs were pooled from multiple independent preparations and divided between treatment groups (**Figure 3A**). NGS analysis revealed that PRIME-VLP consistently outperformed the standard approach across all targets, with fold improvements ranging from 1.5-fold (*PCNA* +2 AA>CG) to 2.8-fold (HEK3 +1 T del) (**Figure 3B and 3C**).

**Figure 3.**
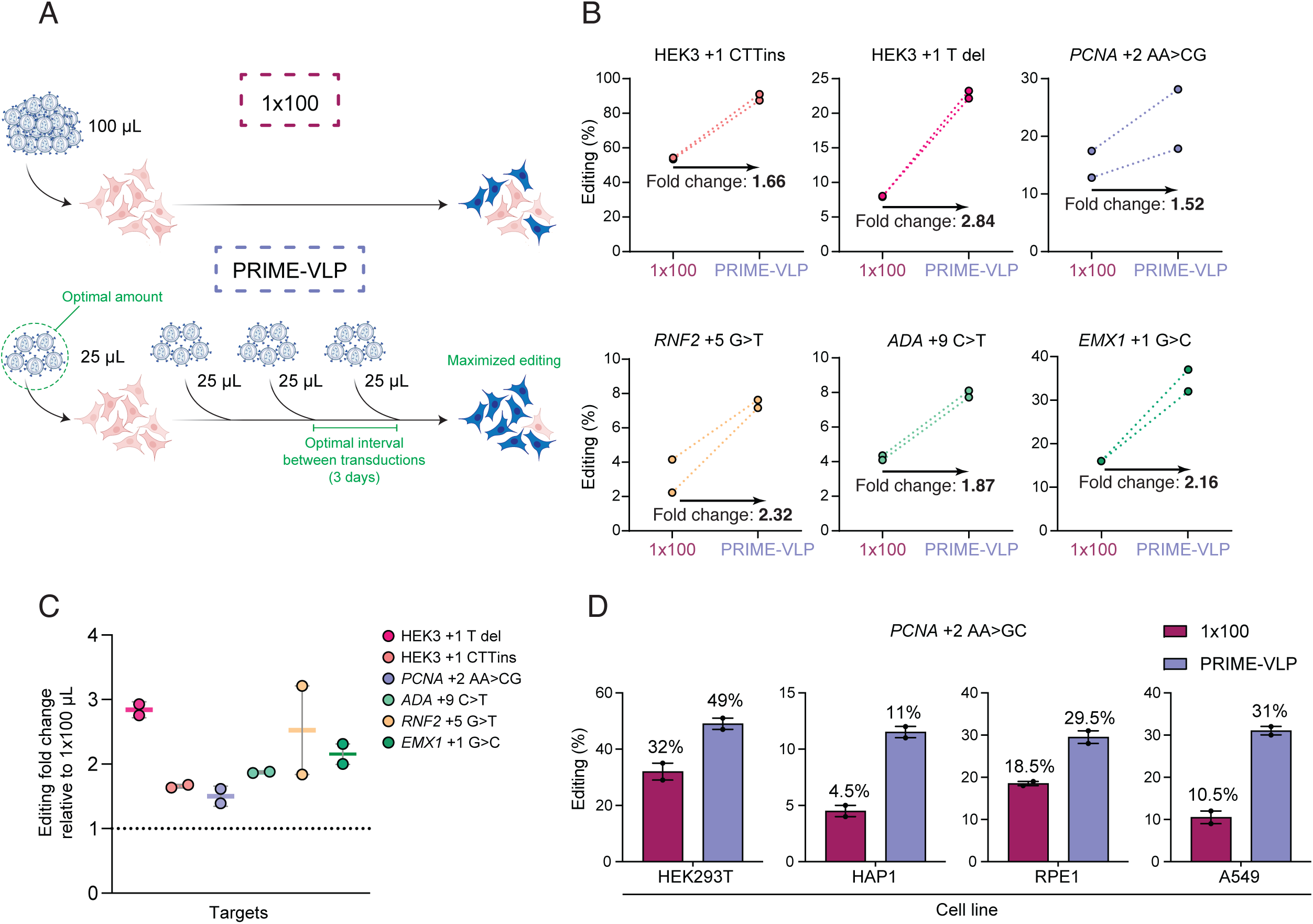
PRIME-VLP improves prime editing efficiency through compound eVLP transductions across diverse targets and cell types. **(A)** Schematic comparison of delivery strategies. Top: single high-dose transduction (1x100 μL) delivering all particles simultaneously. Bottom: PRIME-VLP method using four sequential sub-saturating doses (4x25 μL) delivered at optimal 3-day intervals to maximize editing while avoiding saturation effects. **(B)** Head-to-head comparison of editing efficiency between single high-dose (1x100, red) and PRIME-VLP (purple) delivery across six endogenous genomic targets representing diverse edit types. PRIME-VLP consistently achieves superior editing with fold improvements ranging from 1.52 to 2.84. Individual data points represent independent experiments with connecting lines showing paired comparisons. **(C)** Scatter plot showing fold-change improvements achieved by PRIME-VLP relative to single high-dose treatment (dotted line at y = 1 indicates equivalent performance). Each target is color-coded and shows consistent improvements above baseline, with most targets achieving 1.5-3-fold enhancement. **(D)** Comparison of single high-dose (1x100, red) versus PRIME-VLP (purple) delivery across four human cell lines (HEK293T, HAP1, RPE-1, A549) using the PCNA +2 AA>GC target. PRIME-VLP demonstrates consistent improvements across all tested cell types, with percentages above bars indicating absolute editing efficiencies. Data represent mean ± s.e.m. from n = 2 independent experiments.

Importantly, this improvement was observed across targets with widely varying baseline efficiencies, indicating that PRIME-VLP enhances editing regardless of initial efficiency. To assess the broader applicability of PRIME-VLP, we compared PRIME-VLP against single-dose delivery in three additional human cell lines: HAP1, a near-haploid cell line; A549, a lung carcinoma cell line; and RPE-1, a non-transformed retinal pigment epithelial cell line. We employed the *PCNA* +2 AA>CG pegRNA for this analysis, as it demonstrated the most modest improvement (1.5-fold) among our initial targets (**Figure 3B**), thus providing a conservative assessment of PRIME-VLP performance. Enhanced editing efficiency was observed across all cell lines using PRIME-VLP (**Figure 3D**). In HEK293T cells, editing increased from 32% to 49%, while similar improvements were achieved in HAP1 (4.5% to 11.5%), A549 (10.5% to 31%), and RPE-1 (18.5% to 29.5%). Due to the challenges of performing prime editing in primary human T cells, primarily because mismatch repair competency reduces prime editing efficiencies^19,20,56^, we tested whether PRIME-VLP could be applied to primary cells using base editing instead. T cells were edited with ABE-eVLPs using either single high-dose transduction or PRIME-VLP at multiple endogenous loci. While single high-dose eVLP delivery yielded robust editing across all targets (average: 31.9%), PRIME-VLP further enhanced editing, achieving an average of 44.6%, a 1.4-fold increase (**Figure S3A**). These results demonstrate that PRIME-VLP enhances editing across diverse genomic loci, cell lines, including primary T cells, and genome editing platforms, establishing its versatility for precision genome editing applications.

### Off-target analysis demonstrates enhanced on-target precision of PRIME-VLP

Enhanced editing efficiency could potentially exacerbate off-target activity through prolonged or repeated editor exposure. To address this concern, we conducted an off-target analysis in HEK293T cells using two promiscuous gRNAs with established off-target profiles^13^ and compared PRIME-VLP to single high-dose transduction. As a positive control, we delivered Cas9 nuclease via plasmid transfection or eVLPs and measured editing at both on-target sites and eight established off-target sites using NGS. Plasmid delivery of Cas9 resulted in successful on-target indel generation (34.4% at HEK3, 16% at HEK4), accompanied by substantial off-target editing at all sites, with one off-target site for HEK4 generating more indels (off-target site 3: 59.4%) than the on-target site (**Figure 4A**). Cas9-eVLP delivery dramatically enhanced on-target editing efficiencies (up to 98.4% for HEK3, 99.5% for HEK4) compared to plasmid transfection, while retaining activity at all off-target sites, confirming both the robust editing capacity of eVLPs and the promiscuous nature of these gRNAs (**Figure 4A**). These data validated that our selected gRNAs exhibit robust, detectable off-target activity, providing a sensitive system for evaluating PRIME-VLP specificity.

**Figure 4.**
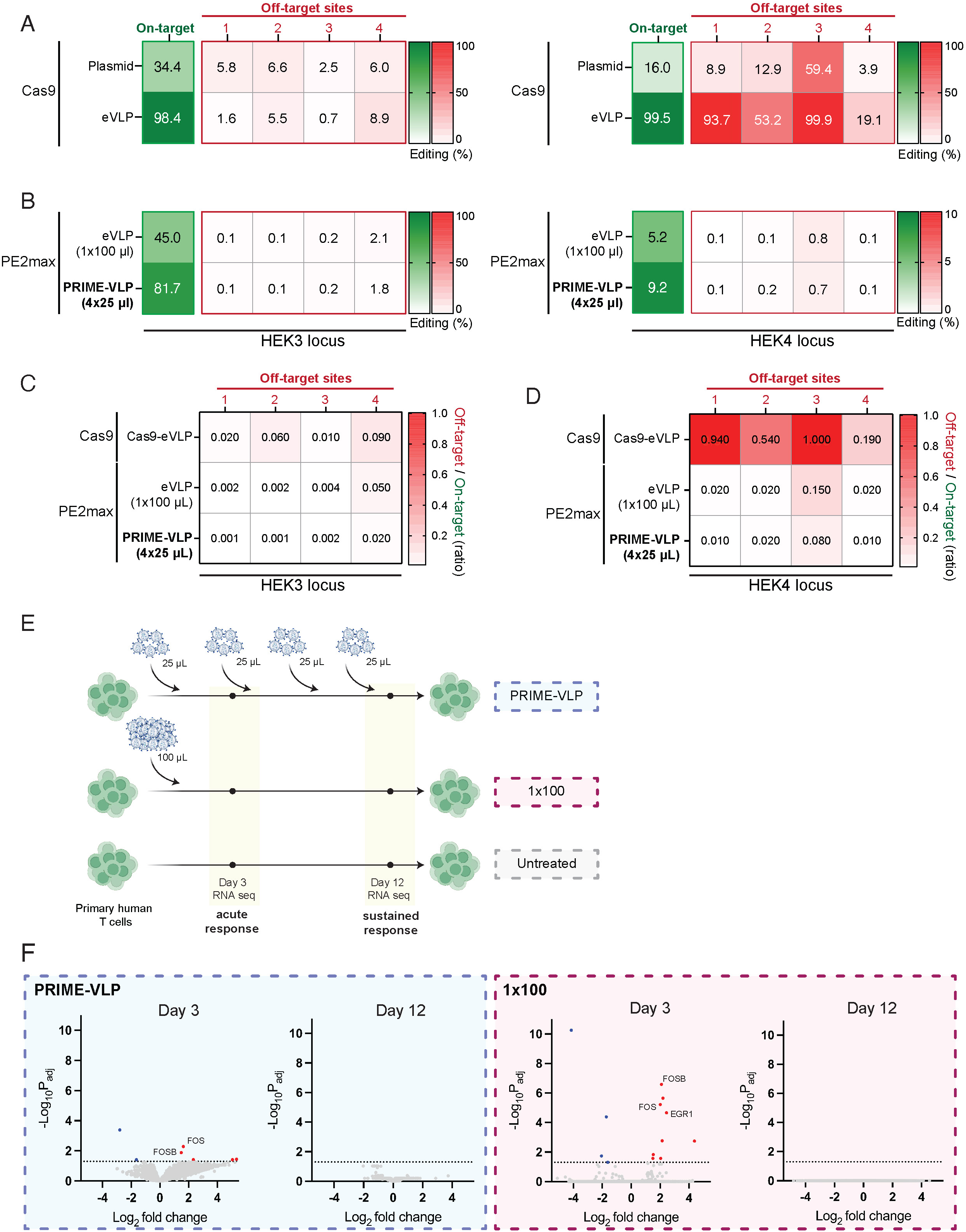
PRIME-VLP maintains editing specificity without transcriptome perturbation. **(A)** Off-target editing analysis using promiscuous HEK3 and HEK4 gRNAs with established off-target profiles. Heat maps compare Cas9 nuclease delivery via plasmid transfection versus eVLPs across on-target sites and four characterized off-target sites. Values represent absolute editing percentages, with green indicating on-target editing and red indicating off-target editing sites. eVLP delivery achieves superior on-target editing while maintaining detectable off-target activity, validating the sensitivity of this assay system. **(B)** Prime editing specificity comparison between single high-dose (1x100 μL) and PRIME-VLP (4x25 μL) delivery using the same promiscuous gRNAs. Heat maps show that PRIME-VLP achieves substantially higher on-target editing (45.0% vs 81.7% for HEK3; 5.2% vs 9.2% for HEK4) without proportional increases in off-target editing across any of the four characterized sites. **(C and D)** Off-target to on-target editing ratios for HEK3 (C) and HEK4 (D) loci demonstrating improved specificity profiles. Heat maps display the ratio of off-target editing relative to on-target editing, with lower values (green) indicating better specificity. PRIME-VLP maintains or improves specificity ratios compared to single high-dose delivery across all off-target sites tested. **(E)** Experimental design for transcriptome analysis in primary human T cells from four healthy donors. Timeline shows comparison of three conditions: PRIME-VLP (4x25 μL, top), single high-dose (1x100 μL, middle), and untreated controls (bottom). RNA sequencing was performed at two timepoints: day 3 (acute cellular response) and day 12 (sustained response assessment). **(F)** Volcano plots of differential gene expression analysis comparing PRIME-VLP and single high dose conditions. PRIME-VLP (left) induces fewer and less pronounced transcriptional changes compared to single high-dose delivery (right) at day 3. By day 12, both conditions return to baseline transcriptional states. Significantly upregulated genes (red) and downregulated genes (blue) are highlighted, with specific genes labeled (FOSB, FOS, EGR1). Dotted horizontal line indicates significance threshold (adjusted p-value < 0.05).

We next compared on- and off-target activities between PRIME-VLP (4 x 25 μL) and the same number of particles in a single high-dose PE2max-eVLP transduction (1 x 100 μL) (**Figure 4B**). Consistent with our previous observations (**Figure 3**), PRIME-VLP achieved significantly higher on-target editing with a nearly 2-fold increase for both pegRNAs (45% to 81.7% for HEK3 and 5.2% to 9.2% for HEK4) (**Figure 4B**). Critically, this substantial increase in on-target editing was not accompanied by detectable increases in off-target editing at any of the off-target sites examined, with the off-target to on-target editing ratios being maintained or improved at both HEK3 (**Figure 4C**) and HEK4 (**Figure 4D**). These results demonstrate that PRIME-VLP enhances on-target editing efficiency while maintaining favorable off-target profiles, indicating its improved editing specificity.

### Transcriptome and cellular viability analysis demonstrate minimal cellular perturbation

Beyond off-target editing concerns, repeated eVLP transductions could potentially perturb cellular physiology or gene expression. We therefore performed transcriptome-wide analysis and cellular viability assessments. Using bulk RNA sequencing, we compared transcriptional profiles of primary human T cells from four healthy donors treated with a single high-dose eVLP (100 μL) or the same number of particles using PRIME-VLP (4 x 25 μL), relative to untreated control cells (**Figure 4E**). To capture both acute and sustained transcriptional responses, cells were harvested and RNA sequenced at two critical timepoints: three days after the initial transduction and after the final transduction (**Figure 4E and S4A**). This experimental design enables the detection of both potential immediate cellular stress responses and long-term transcriptional dysregulation.

Differential expression analysis revealed modest transcriptional perturbations on day 3, with single high-dose transduction inducing a stronger transcriptional response than PRIME-VLP (**Figure 4F**). This differential response is likely due to the four-fold higher particle exposure in the high-dose group at this early point, further justifying the importance of using optimized sub-saturating doses. The single high-dose treatment upregulated multiple genes, whereas PRIME-VLP induced more limited changes (**Figure 4F**). Importantly, these transcriptional changes were transient. By day 12, primary T cells had returned to their baseline transcriptional state (**Figure 4F**), indicating that neither delivery approach caused lasting transcriptional dysregulation. To identify pathway-level perturbations, we performed gene set enrichment (fGSEA) analysis^57^ focusing on pathways critical for T cell function, including activation, apoptosis and exhaustion. Notably, no pathways showed significant enrichment in either condition following the initial response (**Figure S4B**). Principal component analysis revealed that samples clustered primarily by timepoint and donor, with delivery method having minimal influence, indicating that temporal dynamics (92%) and inter-donor variability (5%) had substantially greater impact on gene expression than either eVLP delivery approaches (**Figure S4C and S4D**).

To assess whether eVLP delivery affected cell viability, we measured cell viability in both primary human T cells and HEK293T cells treated with a single high-dose transduction (1 x 100 μL) or PRIME-VLP (4 x 25 μL) (**Figure S5A**). Cell viability was assessed in T cells on day 3, coinciding with the observed transcriptional response peak. No significant differences in cell viability were observed compared to untreated controls (**Figure S5B**). A transient reduction in T cell expansion was observed between days 3 and 6 in eVLP-treated samples; however, this effect was equivalent between PRIME-VLP and high-dose conditions and did not persist beyond day 6 (**Figure S5C**). Similarly, HEK293T cells monitored at regular intervals showed that neither approach affected cell viability relative to untreated controls **(Figure S5C)**.

Collectively, these analyses demonstrate that PRIME-VLP maintains cellular homeostasis across multiple cellular endpoints. Despite repeated eVLP exposure, PRIME-VLP induces only transient, mild transcriptional responses that resolve completely without pathway-level disruption or sustained viability effects. These findings establish that the enhanced editing efficiency of PRIME-VLP is achieved without compromising cellular homeostasis.

### Decoupling editor and (pe)gRNA delivery enables high-throughput prime editing screens using eVLPs

Having demonstrated that PRIME-VLP enhances prime editing efficiency without compromising specificity or cellular fitness, we sought to adapt this approach to high-throughput prime editing screens, where low editing efficiencies represent a major technical barrier^27,29,30,58^. Previous work identified pegRNA packaging as a key limitation in eVLP-based delivery systems^41,46^. Consequently, aptamer-based approaches, such as PP7^46^ and MS2^41^, have been developed to enhance pegRNA packaging efficiency, allowing more pegRNAs to be loaded per particle. However, we decided to pursue a fundamentally opposite approach by using pegRNA-free particles that decouple the editor and pegRNA delivery entirely. We hypothesized that eVLPs could either deliver pegRNA-free editors or complexed with a non-targeting pegRNA that could be functionally exchanged by a lentivirally-delivered pegRNA library, similar to what was previously demonstrated with gRNAs and Cas9 RNP electroporation^59^.

To test the pegRNA swapping strategy, we produced PE2max-eVLPs containing either targeting pegRNAs (positive control), non-targeting pegRNAs, or no pegRNAs. We hypothesized that eVLPs lacking targeting pegRNAs could still achieve functional editing when targeting pegRNAs are delivered independently via lentivirus (**Figure 5A**). We evaluated this approach using three distinct pegRNAs introducing a variety of mutation types (HEK3 +1 CTTins, HEK3 +1 del, *PCNA* +2 AA>CG). Our results demonstrated successful pegRNA exchange across all tested conditions (**Figure 5B**). eVLPs loaded with targeting pegRNAs achieved an average editing efficiency of 73.6% (**Figure 5B**) across all three edit types (range: 55.9-97.3%, (**Figure S6A**)). Importantly, editors delivered with non-targeting pegRNAs successfully swapped with lentivirally-delivered targeting pegRNAs, achieving editing efficiencies averaging 68.7% (**Figure 5B**) across all three edits (range: 50.8-91.4%, (**Figure S6A**)), representing only a modest reduction compared to the positive control. Notably, pegRNA-free eVLPs retained substantial editing capacity across all edits, indicating that while non-targeting pegRNAs enhance prime editor stability and editing efficiency, they are not strictly required for functional editing (**Figure 5B and S6A**).

**Figure 5.**
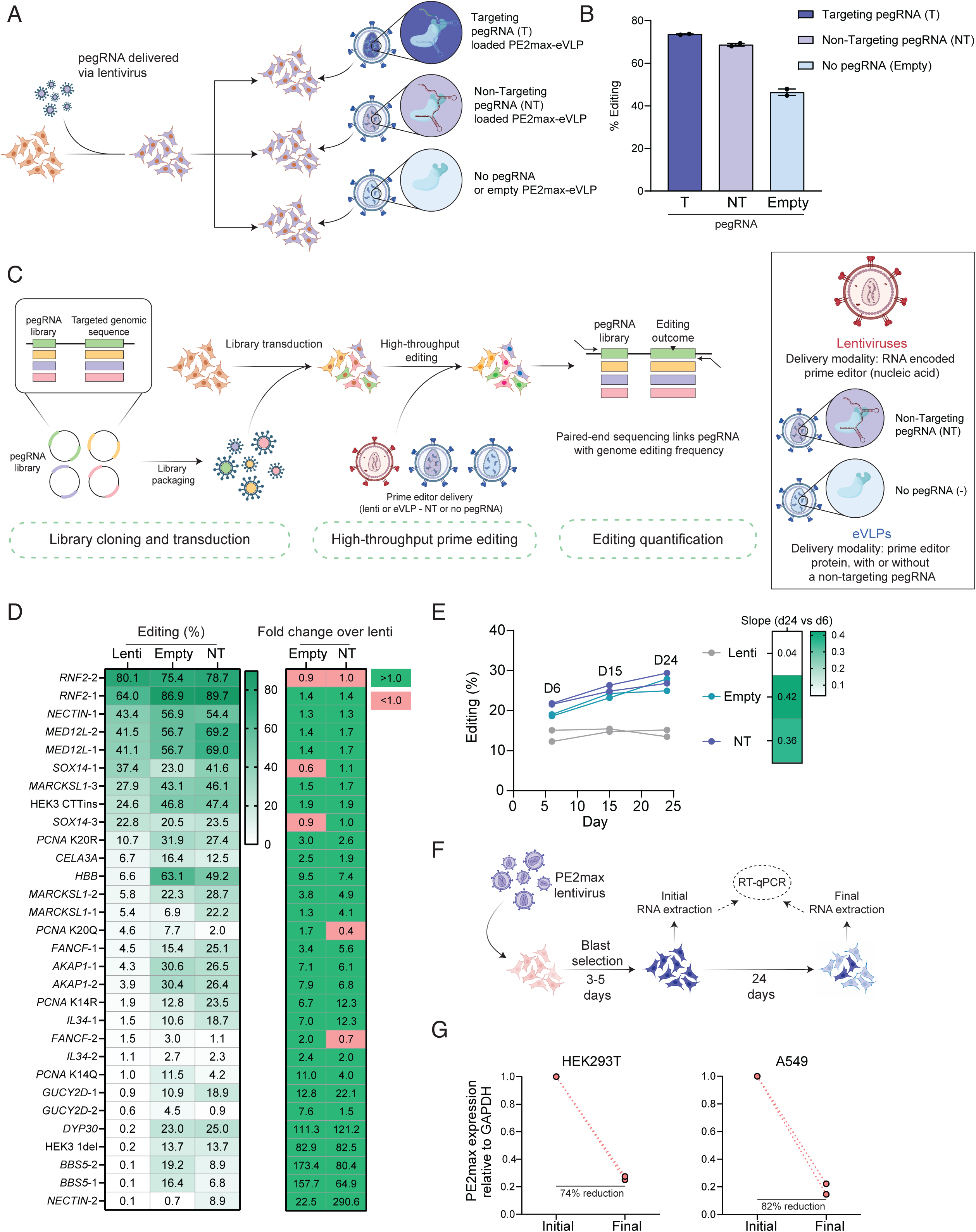
Decoupling editor and pegRNA delivery enables high-throughput prime editing screens. **(A)** Schematic of the decoupled delivery strategy. pegRNAs are delivered via lentiviral transduction (left), while prime editors are delivered separately using eVLPs containing three different configurations: targeting pegRNAs (T, dark blue), non-targeting pegRNAs (NT, blue), or no pegRNAs/empty eVLPs (-, pale blue). This approach separates library delivery from editor delivery to overcome lentiviral expression limitations. **(B)** Validation of pegRNA exchange functionality across three edit types (HEK3 +1 CTTins, HEK3 +1 T del, *PCNA* +2 AA>CG). PE2max-eVLPs with targeting pegRNAs (T) achieved highest editing efficiency (73.6% average), while non-targeting pegRNAs (NT) showed comparable performance (68.7% average), and empty eVLPs (-) retained substantial activity (43.8% average), confirming successful editor-pegRNA decoupling. Data represent mean ± s.e.m. **(C)** Miniature sensor library screening workflow. The mini sensor library contains each pegRNA paired with its corresponding target sequence within the same construct. Following library transduction and prime editing, paired-end sequencing quantifies both pegRNA abundance and editing outcomes at the linked target sequences, enabling high-throughput assessment of editing efficiency. **(D)** Performance comparison in the miniature sensor screen across all 30 targets. Left: Individual target editing efficiencies showing PRIME-VLP conditions (Empty and NT) consistently outperform lentiviral PE2max delivery (Lenti) with approximately 2-fold higher mean editing efficiency (Empty: 29.1%, NT: 27.3%) compared to lentiviral delivery (14.8%). Right: Heat map of fold-change improvements for Empty and NT conditions relative to lentiviral delivery, with green indicating superior eVLP performance. **(E)** Temporal analysis of editing accumulation over 24 days. Line graph shows sustained editing accumulation for PRIME-VLP conditions (Empty and NT) with positive slopes (0.42 and 0.36, respectively), while lentiviral delivery plateaus after day 6 with minimal further improvement (slope = 0.04). This demonstrates the superior sustainability of protein-based editor delivery. **(F)** Experimental design for PE2max expression stability analysis. Cells are transduced with PE2max lentivirus, selected with blasticidin, and analyzed by RT-qPCR at initial timepoint (post-selection) and after 24 days to assess transgene silencing over time. **(G)** Quantitative RT-qPCR analysis of PE2max transcript levels relative to *GAPDH* housekeeping gene. Significant reductions in PE2max expression are observed in both HEK293T (74% reduction) and A549 (82% reduction) cell lines over 24 days, explaining the superior sustained performance of protein-based eVLP delivery over lentiviral systems. Individual biological replicates are shown with connecting lines.

To assess the generalizability of (pe)gRNA decoupling across different CRISPR systems, we extended these analyses to base editing. We tested ABE-eVLPs targeting three independent loci (HEK3, *RASA2*, and *FANCF*) using the same decoupling strategy (**Figure S6B**). Similar patterns were observed with ABE-VLPs, with particles containing targeting gRNAs achieving an average of 95.4% editing efficiency, those with non-targeting gRNAs reaching 76.0% efficiency, and gRNA-free particles achieving 68.0% efficiency (**Figure S6C and S6D**). These results demonstrate that the eVLP decoupling strategy is broadly applicable across different CRISPR-based editing platforms with a limited loss of activity.

Altogether, these findings establish that editors and (pe)gRNAs can be delivered independently while maintaining high editing efficiency, providing a robust foundation for high-throughput screening applications using eVLPs. This decoupling strategy is critical for screening applications, as it enables eVLP-based delivery of editors using (pe)gRNA-free particles while introducing the library via lentiviruses.

### PRIME-VLP enhances high-throughput prime editing screens

To evaluate whether PRIME-VLP enhances pooled prime editing screen efficiency, we constructed a miniature (mini) pegRNA sensor library in which each pegRNA is paired with its surrogate genomic target sequence, enabling high-throughput quantification of editing efficiency by NGS (**Figure 5C**). We optimized eVLP dosing parameters using non-targeting pegRNA and pegRNA-free eVLPs specifically for pooled screening applications across diverse cell lines. HEK293T, RPE-1, K562 and HCT116 cells were transduced with our mini sensor library at low MOI (MOI = 0.3) to ensure one integration per cell. Following library selection, cells were treated with increasing amounts (25-100 μL) of pegRNA-free PE2max-eVLPs. NGS analysis revealed a consistent dose-response relationship across all cell lines, with a small increase in editing efficiency between 25 μL and 50 μL before reaching saturation at higher doses (**Figure S7A**). These titrations identified 50 μL as the optimal dose for PRIME-VLP using decoupled pegRNAs. We also compared our PE2max-eVLPs with the recently reported eVLPv2.3 system^41^ across four different cell lines and found highly comparable performance, with v2.3 achieving a modest 1.09-fold improvement over PE2max-eVLPs (**Figure S7B and S7C**). Based on these optimizations, we adopted PE2max-eVLPv2.3 with 50 μL dosing for subsequent pooled screening experiments.

To benchmark PRIME-VLP performance against established methodologies, we conducted direct comparisons between eVLP-mediated editor delivery and conventional lentiviral delivery systems in a pooled prime editing sensor screen. HEK293T cells were transduced with the pegRNA library and subsequently treated with prime editors delivered via lentivirus (Lenti), eVLPs loaded with a non-targeting pegRNA (NT), or pegRNA-free eVLPs (Empty). Over a 24-day screening experiment comprising eight PRIME-VLP transductions, we harvested cells at multiple timepoints (days 6, 15 and 24) to assess temporal editing dynamics. Lentiviral delivery of PE2max exhibited highly variable editing efficiency ranging from 0.1% to 80.1% across different loci, with a mean of 14.8% (**Figure 5D**). In striking contrast, PRIME-VLP consistently outperformed lentivirus at nearly every locus, with both Empty and NT conditions achieving average editing efficiencies of 29.1% and 27.3%, respectively, representing approximately a 2-fold improvement over lentiviral delivery (**Figure 5D**). These results highlight the potential of PRIME-VLP as a scalable high-throughput editing platform, and demonstrate that robust prime editing can be achieved without pre-loading pegRNAs into eVLPs.

To understand the mechanism underlying PRIME-VLP’s superior performance, we analyzed the temporal dynamics of editing accumulation. Both PRIME-VLP approaches (Empty and NT) demonstrated robust initial editing (day 6), followed by sustained accumulation throughout the experiment (slope = 0.42 and 0.36) (**Figure 5E**). In contrast, lentiviral delivery of PE2max exhibited minimal editing accumulation beyond day 6 (slope =0.04), suggesting a plateau in editing efficiency after the initial week (**Figure 5E**). This temporal difference suggested that lentiviral PE2max expression might decline over time, potentially due to transgene silencing^33,34,60^.

To test this hypothesis, we measured PE2max expression over time in multiple cell lines. We transduced HEK293T, A549, HAP1, HCT116, and K562 cells with PE2max lentivirus, selected for stable integration, and quantified PE2max expression relative to *GAPDH* by RT-qPCR at day 0 (post-selection) and day 24 (**Figure 5F**). The results confirmed substantial transgene silencing across all cell lines, with PE2max transcript levels declining by 74% in HEK293T cells, 82% in A549 cells (**Figure 5G**), and 27-53% in HAP1, HCT116 and K562 (**Figure S8A**). These experiments demonstrate that diminished PE2max expression impairs editing efficiency, highlighting transgene silencing as a major limitation of lentiviral systems that is circumvented by RNP-based eVLP delivery.

### PRIME-VLP enables scalable high-throughput prime editing screens and enables functional variant discovery

To demonstrate the scalability and practical utility of PRIME-VLP in large-scale functional genomics applications, we evaluated its performance using a biologically relevant, higher complexity screening platform. We designed a 6,000 pegRNA library targeting *TP53* that includes functionally validated variants with known phenotypic effects^29^, and employs positive selection to identify pathogenic *TP53* mutations that confer resistance to Nutlin-3^61^.

We first assessed baseline editing efficiency without phenotypic selection to benchmark PRIME-VLP performance against conventional lentiviral PE2max delivery in a large-scale pooled format (**Figure 6A**). HEK293T cells, which lack wild-type *TP53* expression, were transduced with the 6,000-pegRNA library at low multiplicity of infection (MOI) (MOI = 0.3), followed by PRIME-VLP treatment using PE2max-eVLPv2.3 loaded with non-targeting pegRNAs over a 24-day period (**Figure 6A**). NGS of the pegRNA and surrogate sensor sites recovered 5,784 pegRNAs (96.4% library coverage) and revealed that PRIME-VLP achieved consistently high editing efficiencies across all functional domains of p53 (**Figure S9A**). Remarkably, 82.1% of pegRNAs exhibited editing above 10%, compared to only 44.2% of pegRNAs achieving this threshold with lentiviral delivery (**Figure 6B**), representing a 1.9-fold improvement in the proportion of functionally active pegRNAs. PRIME-VLP editing efficiencies were consistently high across pegRNAs, with a mean of 44.1%, a 2.8-fold increase over lentiviral delivery (15.9%) (**Figure 6C**). Notably, both PRIME-VLP and lentiviral replicates demonstrated excellent reproducibility with mean editing differences of only 0.5% and 0.7% between replicates, respectively, thereby validating the robustness of the screens and enabling accurate comparison between delivery methods (**Figure 6C**). On an individual pegRNA level, 4,319 pegRNAs showed substantial improvements (>10% increase in absolute editing efficiency) over lentivirus, while only 1,053 showed modest gains (0-10% increase), and only 412 (7.1%) showed improvements in the lentivirus condition (**Figure 6D**). We next assessed the pegRNA-level correlation between experimental replicates across the library. While PRIME-VLP showed strong concordance for individual pegRNAs between replicates (R² = 0.82), indicating consistent editing outcomes for each target, the lentiviral condition exhibited poor pegRNA-level reproducibility (R² = 0.2) (**Figure S9B**). This difference reflects the inherent variability in lentiviral systems, where random transgene integration sites lead to stochastic and mosaic expression patterns and silencing across cell populations, resulting in inconsistent editing outcomes for individual pegRNAs despite similar overall mean performance. To confirm that the superior performance of PRIME-VLP was not simply due to technical limitations of the lentiviral replicates, we compared the best-performing lentiviral replicate for each pegRNA against the average PRIME-VLP performance. Even under these conditions that favored the lentiviral approach, PRIME-VLP outperformed lentiviral PE2max by 1.9-fold (**Figure S9E**). Additionally, PRIME-VLP achieved more uniform editing efficiency across the entire pegRNA library, particularly benefiting lower-efficiency pegRNAs that would otherwise perform too poorly for reliable screening applications (**Figure S9C**). Importantly, the reduced editing efficiency observed with lentiviral delivery was not due to poor library representation or low sequencing quality, as both conditions showed comparable pegRNA coverage and read depth (**Figure S9D**). Together, these results establish PRIME-VLP as a robust platform for high-throughput prime editing screens, enabling both efficient and reproducible editing across thousands of targets.

**Figure 6.**
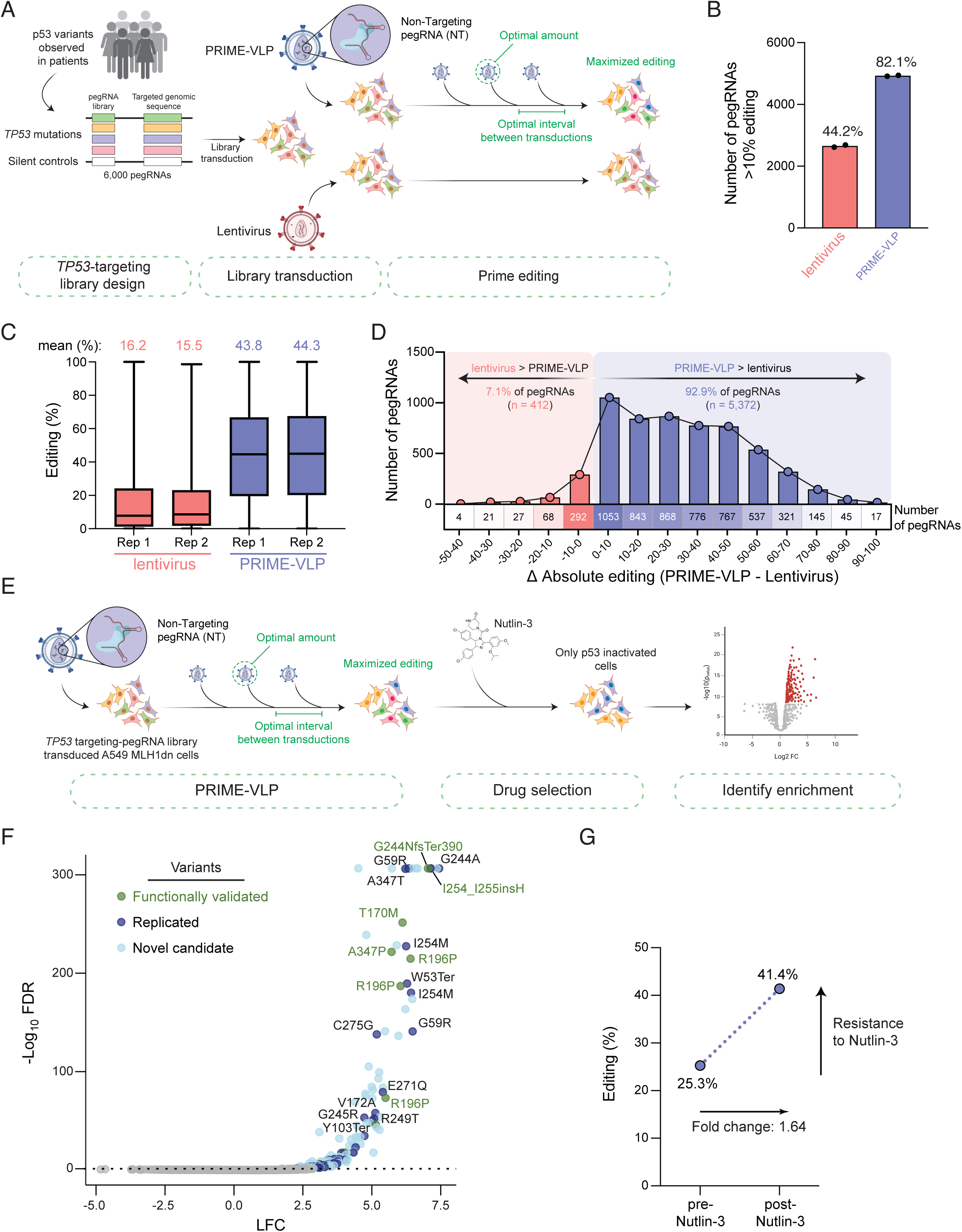
PRIME-VLP enables large-scale functional screening. **(A)** Experimental design for the 6,000-pegRNA *TP53* sensor library screen. The library targets p53 functional domains and includes known patient variants alongside silent controls. HEK293T cells are transduced with the pegRNA library via lentivirus, followed by prime editor delivery using either lentiviral PE2max (bottom path) or PRIME-VLP with non-targeting pegRNA-loaded eVLPs (top path) delivered at optimal intervals over 24 days. **(B)** Comparison of functionally active pegRNAs achieving >10% editing efficiency. PRIME-VLP achieves 82.1% active pegRNAs compared to 44.2% with lentiviral delivery, representing a 1.9-fold improvement in functional library coverage. Data represent mean ± s.e.m. from n = 2 independent biological replicates. **(C)** Distribution of editing efficiencies across biological replicates. Box plots demonstrate that PRIME-VLP achieves 2.8-fold higher mean editing efficiency (44.1%) compared to lentiviral delivery (15.9%). PRIME-VLP also shows superior reproducibility between replicates (mean difference: 0.5%) compared to lentiviral delivery (0.7%). Box plots display median, quartiles, and range across all pegRNAs for n = 2 biological replicates per condition. **(D)** pegRNA-level performance comparison across the entire library. Histogram shows the distribution of absolute editing differences (PRIME-VLP minus lentivirus) binned by percentage point improvements. 92.9% of pegRNAs (5,372 out of 5,784) demonstrate improved editing with PRIME-VLP, while only 7.1% (412 pegRNAs) show better performance with lentiviral delivery, highlighting the broad superiority of the PRIME-VLP approach. **(E)** Functional screening workflow for *TP53* loss-of-function variant identification. A549 cells (wild-type *TP53*) are transduced with the *TP53* pegRNA library and subjected to PRIME-VLP treatment over 24 days. Cells are then treated with Nutlin-3 to select for variants that confer resistance through p53 pathway disruption. The volcano plot illustrates the selection outcome, with resistant variants enriched in the surviving population. **(F)** MAGeCK analysis results from Nutlin-3 selection screen. Volcano plot displays statistical significance (-log10 FDR) versus effect size (log fold change, LFC) for pegRNA enrichment after 11 days of Nutlin-3 treatment. 181 significantly enriched pegRNAs were identified (FDR < 0.05). Color coding indicates: overlapping hits with previous studies (dark blue), validated overlapping hits (green), and unique hits identified in this study (light blue). Key variants are labeled, including hotspot mutations and novel candidates. **(G)** Validation of functional selection through editing efficiency analysis. Scatter plot shows editing efficiency for the 62 overlapping hits before (pre-) and after (post-) Nutlin-3 selection, demonstrating successful enrichment of resistance-conferring mutations. Mean editing efficiency increased from 25.3% to 41.4% (1.64-fold change), confirming that cells harboring *TP53* loss-of-function edits preferentially survived Nutlin-3 treatment.

Having benchmarked PRIME-VLP performance in a high-throughput context, we next evaluated its utility for functional screening applications. We employed the 6,000-pegRNA *TP53* library to perform a positive selection screen in A549, a lung adenocarcinoma cell line with wild-type *TP53* and intact p53 signalling. Cells were transduced with the library at a low MOI and subjected to compound PRIME-VLP transductions every 3 days over a 24-day period using PE2max-eVLPv2.3 loaded with a non-targeting pegRNA (**Figure 6E**). Following editing, cells were treated with Nutlin-3, and genomic DNA was harvested at day 11 post-treatment for NGS to assess pegRNA frequencies and sensor editing outcomes. Although prime editing at the surrogate sites in A549 cells was lower than in HEK293T cells, which is consistent with our previous observations (**Figure 3D**), sensor editing was still robust across p53 domains (**Figure S10A**). Following Nutlin-3 treatment, we observed a notable 1.9-fold increase in editing efficiency across the library (**Figure S10A**), consistent with the selective enrichment of cells harbouring *TP53* loss-of-function mutations that confer resistance to p53-mediated growth arrest. Using the sensor library enabled us to calibrate our dataset by filtering for active pegRNAs, as done previously^29,62^. After filtering for active pegRNAs (>10% editing), MAGeCK analysis of pegRNA abundance at day 11 post-Nutlin treatment identified 181 significantly enriched pegRNAs, successfully capturing 62 replicated variants^29^ and identifying novel candidate *TP53* mutations (**Figure 6F**). These overlapping hits had an average editing efficiency of 25.3% before Nutlin-3 treatment, which increased to 41.4% after treatment, highlighting their resistance to Nutlin-3 (**Figure 6G**). On an individual pegRNA level, 50 out of 62 overlapping hits showed substantial enrichment in editing efficiency, reflecting the selective survival and proliferation of edited cells that resist Nutlin-3-induced growth inhibition, while unedited cells undergo cell cycle arrest (**Figure S10B**). Importantly, our PRIME-VLP screen recapitulated 11 functionally validated p53 variants^29^, including several hotspot mutations targeting R248 (**Figure S10C**). All of these variants showed substantial increases in editing frequency post-selection; for example, G244NfsTer390 rose from 20.2% to 58.3%, and T170M from 17.9% to 41%, demonstrating resistance to Nutlin treatment (**Figure S10D**). The screen also identified variants affecting the same amino acid residues as functionally validated hits, such as G244A, I254M and A347T (**Figure 6F**), suggesting similar mechanisms of p53 inactivation. These findings demonstrate that PRIME-VLP enables robust, high-fidelity pooled prime editing screens in a biologically relevant context, identifying functional variants without requiring stable transgenic expression of a prime editor and providing an effective alternative to conventional lentiviral delivery for functional CRISPR screens.

## DISCUSSION

Advances in sequencing technologies have led to the identification of an increasing number of genetic variants of unknown significance, necessitating scalable approaches for their functional characterization^1–5^. Prime editing offers a powerful means to precisely introduce these variants without requiring double-strand breaks or donor templates, making it uniquely suited for such applications^13,15^. However, its adoption, particularly for high-throughput screening, has been limited by suboptimal editing efficiencies. While recent efforts have focused on improving prime editor design^20–23^ and pegRNA architecture^16–18^, alternative delivery strategies remain underexplored. Here, we introduce PRIME-VLP, an eVLP-based delivery strategy that utilizes compound delivery of multiple temporally spaced, sub-saturating particle doses to achieve consistent, high-efficiency prime editing across diverse genomic targets, variants, cell types, and throughput.

Our kinetics analysis revealed that eVLP-mediated prime editing reaches maximum editing within three days post-transduction, indicating a narrow window of peak RNP activity. This is consistent with prior studies of DNA repair kinetics showing that Cas9 activity plateaus around 60 hours post-induction^63^. The dose-response analysis further revealed saturation effects at higher eVLP concentrations, where increasing particle numbers yielded diminishing returns in editing efficiency. This plateau likely reflects intracellular constraints such as target site accessibility, chromatin dynamics, or competition among RNPs. Pre-assembled RNPs likely bind available targets rapidly upon entry, while excess RNPs are degraded before engaging new sites, an observation supported by the reported Cas9 half-life of 6–8 hours of delivery^63^. These kinetic and dose-response insights provided the rational foundation for developing PRIME-VLP, as they indicated that multiple sub-saturating doses delivered at optimal intervals could maintain steady-state editor levels while avoiding saturation-induced limitations.

By conducting sequential eVLP transductions in HEK293T cells and human primary T cells, we observed that cells remained permissive to eVLP uptake and editing across multiple rounds, with secondary transductions producing comparable or additional editing between previously transduced and naïve cells (**Figure 2D and S2D**). The absence of viral nucleic acids in eVLPs likely contributes to this sustained transduction capacity by avoiding activation of cytosolic nucleic acid sensors, including cGAS-STING and RIG-I/MDA5^55,64,65^. Our results demonstrate that sequential delivery of optimized eVLP doses prevents saturation effects and achieves superior cumulative editing efficiency compared to single high-dose treatments (**Figure 3)**. However, when the same genomic locus was targeted across multiple rounds of transduction, editing levels increased significantly but did not double as might be expected. This likely reflects both the mathematical constraint imposed by progressive depletion of unedited target sites, as editing depletes unedited targets, and potential locus-specific factors that influence re-editing efficiency. Given the established role of chromatin structure on prime editing accessibility^66,67^, successful editing events may trigger transient chromatin remodelling that creates a refractory period at targeted loci. Additionally, edits that preserve the PAM sequence may be repeatedly targeted, potentially interfering with repair or overwriting prior edits. Nevertheless, PRIME-VLP achieved consistent improvements by exploiting the temporal kinetics of eVLP-delivered RNP complexes to enhance precision genome editing tools.

Off-target and transcriptome analyses demonstrated that PRIME-VLP maintains high specificity while minimizing cellular perturbation. Off-target analysis using promiscuous gRNAs with established off-target profiles revealed that PRIME-VLP enhanced on-target editing while maintaining or improving specificity ratios compared to single high-dose treatments. The temporally distributed RNP delivery in PRIME-VLP may prevent excessive intracellular editor protein accumulation that could drive off-target DNA binding during the target search process. This finding supports the rationale for optimized sub-saturating doses rather than maximal particle delivery. Transcriptomic analysis further supported this finding, showing that PRIME-VLP induced fewer and more transient gene expression changes compared to single high-dose delivery (**Figure 4F**). These stronger initial transcriptional responses induced by the higher dose likely reflect the four-fold higher particle exposure. The upregulation of immediate early genes including *FOS*, *FOSB*, and *EGR1* was consistent with retroviral capsid sensing via TRIM5α, which drives downstream MAP kinase and NF-κB signaling^68^. Additionally, CLMN upregulation may reflect cytoskeletal remodelling during eVLP entry^69^. These findings suggest that excessive particle delivery can trigger cellular stress responses that are minimized with optimized dosing strategies.

PRIME-VLP offers significant advantages for high-throughput prime editing screens by overcoming delivery constraints inherent to existing lentiviral approaches. Current lentiviral systems suffer from packaging limitations for large prime editor constructs, resulting in low functional titers^35,36^, the need for selectable markers^37,53^, and variable expression due to integration site effects^31,32^. We demonstrated that loss of transgene expression^33,34^ further impacts sustained prime editor expression in target cell populations in pooled settings, with PE2max levels declining 27-82% over 24 days across multiple cell lines. This has been somewhat addressed through site-specific transgene integration of prime editors with Cas9-mediated HDR^27^. However, single-cell cloning was required to isolate high-expressing clones, indicating that the integration site is not the only limitation^27^. Moreover, single-cell cloning can lead to the emergence of clone-specific phenotypes, with unique functional characteristics unrelated to the intended genetic modification, often due to genomic instability accumulated during prolonged expansion^70^. Moreover, the process is also labour-intensive, poorly scalable across multiple cell lines, and incompatible with many physiologically relevant models, which are not amenable to single-cell cloning. In contrast, by delivering the pegRNA library via lentivirus and supplying prime editors via eVLPs, we showed that PRIME-VLP circumvents limitations of lentiviral-based editor expression.

When characterizing this pegRNA-free PRIME-VLP system, we discovered that higher eVLP quantities were required in pooled settings compared to targeted genome editing. This likely results from several contributing factors: (1) reduced encounter rates between editor proteins and targeting pegRNAs due to pegRNA instability^18^, (2) inefficient guide swapping in non-targeting conditions leads to the formation of fewer functional RNP complexes loaded with a targeting pegRNA^59^, (3) and decreased editor protein stability in empty eVLPs^71^ may require higher particle numbers to achieve equivalent functional editor concentrations. Interestingly, our data showed that while a non-targeting pegRNA enhances prime editor stability and activity, robust editing is still achievable without it, supporting the use of simplified eVLP formulations in scalable applications. Empty or non-targeting PE2max-eVLPs offer advantages for large-scale screening through their compatibility with diverse library designs and scalable production workflows.

Direct benchmarking against lentiviral delivery demonstrated that PRIME-VLP achieved significantly greater editing, with a 2.8-fold higher average efficiency over a 6,000 pegRNA library (**Figure 6**). Moreover, editing levels across replicates were highly correlated for PRIME-VLP (R² = 0.82), while lentiviral replicates were not (R² = 0.2). This discrepancy reflects the heterogeneous nature of lentiviral expression, in which cells that express high levels of PE2max will likely undergo editing, whereas low-expressing cells will not^32,72^, and these cells are unlikely to contain the same pegRNA in both replicates. These effects are worsened by a progressive decline in transgene expression over time. Across five cell lines, we observed substantial reductions (27-82%) in PE2max transcript levels 24 days after selection (**Figure 5G and S8**). While the mechanism was not directly assessed, this decline may reflect epigenetic silencing, promoter repression or complete transgene loss^33,34^. In contrast, PRIME-VLP delivers the editor as a protein complex, bypassing the need for sustained transcription and resulting in more uniform and consistent editing.

In a large-scale functional screening, PRIME-VLP successfully identified *TP53* loss-of-function mutations conferring resistance to Nutlin-3. Notably, editing for these variants increased substantially following selection, consistent with positive selection. However, the editing frequencies did not reach saturation, as Nutlin-3 primarily induces cell cycle arrest rather than apoptosis^61,73^. Consequently, the 11-day selection window was insufficient to fully outcompete unedited cells, leading to a partial, but substantial, enrichment. Editing in A549 cells was lower than in HEK293Ts, likely reflecting cell-type-specific differences in eVLP uptake, nuclear access, or repair pathway activity^74,75^. Nevertheless, the screen retained sufficient resolution to recover *TP53* variants that promote resistance to Nutlin-3, and identify novel substitutions at key p53 residues, highlighting the robustness of PRIME-VLP for high-throughput prime editing screens. Despite this reduced editing efficiency, employing a sensor-based library enabled filtering for active pegRNAs, significantly enhancing data quality and reducing background noise, as previously demonstrated^29,62^. This approach allowed us to identify loss-of-function variants with high confidence, with 62 replicated variants from previous work^29^, including multiple variants at critical residues such as R248. These results highlight PRIME-VLP’s ability to accurately identify functional variants and identifying novel substitutions at key residues.

Despite the promise of PRIME-VLP, several limitations currently warrant consideration. First, the scalability of eVLP production represents a practical constraint, as total eVLP requirements increase with library complexity and repeated dosing. However, PRIME-VLP offers a critical advantage by using broadly applicable PE2max-eVLPs loaded with non-targeting pegRNAs or empty particles (**Figure 5**), which can be mass-produced independently of specific library designs. Moreover, recent innovations in particle engineering directly address scalability concerns. The eVLPv5 platform incorporates capsid mutations that enhance packaging efficiency and delivery potency by up to 4-fold^42^, while miniEDVs achieve comparable or improved editing with only ∼20% of the original viral composition^47^, thereby reducing particle complexity. These advances will likely reduce the number of particles needed for effective PRIME-VLP and simplify production workflows, helping to overcome current manufacturing constraints. A second limitation of PRIME-VLP is that it relies on VSV-G pseudotyping, which requires *LDLR* expression for cellular entry. Although VSV-G offers broad tropism, LDLR is variably expressed across cell types^76^. For instance, resting or previously activated T cells progressively downregulate LDLR^77,78^, potentially limiting eVLP entry and reducing the benefits of compound dosing. Additionally, differences in endosomal trafficking and LDLR recycling rates between cell types could affect the efficiency of sequential eVLP transductions, explaining why some cell lines benefit more substantially from the PRIME-VLP approach. The varying fold-improvements achieved by PRIME-VLP potentially reflect cell-type-specific differences in LDLR expression levels and receptor recycling kinetics that influence eVLP uptake efficiency (**Figure 3D**). Additionally, whereas lentivirus requires a single integration event, eVLPs must deliver multiple RNP complexes per cell to compensate for protein degradation, making efficient particle uptake particularly critical for achieving high editing levels. However, to overcome this limitation, eVLPs can be readily pseudotyped with alternative glycoproteins to redirect their tropism, highlighting their modularity^40,79,80^. For example, replacing or pairing VSV-G with the baboon endogenous retrovirus (BaEV) envelope has been shown to enhance delivery in low-LDLR-expressing cells^48,81^. Moreover, recent advances employing engineered mutant forms of VSV-G coupled with specific targeting moieties (e.g., single-chain variable fragments) could further broaden the cell-type specificity and improve transduction in hard-to-transduce cell types^43,44,79,82^. A third limitation is that, despite the demonstrated superior efficiency of PRIME-VLP over plasmid transfection (**Figure 1**), single high eVLP dose (**Figure 3**) and lentivirus delivery (**Figures 5 and 6**), the overall editing efficiencies still leave room for improvement. Nevertheless, the modular architecture of eVLPs facilitates rapid incorporation of advances from the genome editing field^14,83^. For instance, identification of a cryptic cleavage site in PE2max prompted adoption of an improved PE2max-eVLPv2.3 construct that resolved this instability. Future enhancements, including optimized prime editor variants, co-delivery of DNA repair modulators (e.g., MLH1dn)^20^, or improved reverse transcriptase enzymes, can be readily integrated into the platform without fundamental redesign. The versatility of PRIME-VLP could also be expanded beyond base and prime editing to other applications that require sustained editor activity. Systems such as epigenome editors^84–86^, rely on prolonged protein presence for effective transcriptional modulation, making them well-suited for compound delivery strategies. Recent successful demonstrations of eVLP-mediated epigenome editor delivery provide additional validation for this expanded utility^87^. Therefore, while current limitations exist, the modularity and adaptability of PRIME-VLP represent significant advantages that allow for rapid integration of recent advancements in genome-editing technologies, particle engineering, and tropism modulation to improve performance and expand the applicability to more diverse cell types. In conclusion, PRIME-VLP introduces a flexible and effective method to overcome key challenges in prime editing delivery for targeted genome engineering and high-throughput screening applications. This work demonstrates how a mechanistic understanding of cellular kinetics can guide the development of more effective genome editing strategies. By leveraging temporally optimized RNP delivery via eVLPs, this approach circumvents limitations associated with lentiviral delivery and enables sustained editing. PRIME-VLP expands the utility of eVLP technology beyond its established *in vivo* therapeutic applications to high-throughput CRISPR-based screening platforms.

## ACKNOWLEDGMENTS

This work was supported by the generous start-up package from the University of Calgary, Cumming School of Medicine, the Arnie Charbonneau Cancer Institute, and the Robson DNA Science Centre to P.B. Grants #20222763 from the Alberta Cellular Therapy and Immune Oncology Initiative and a CIHR grant (508188) to P.B. L.B is supported by the Riddell Centre for Cancer Immunotherapy. J.L was supported by FGS Doctoral Entrance Scholarship, Alberta Graduate Excellence Scholarship and the Phillips Emerging Cancer Scholar Graduate Scholarship.

## AUTHOR CONTRIBUTIONS

J.L, L.B, and J.C have designed and conducted the experiments under the supervision of P.B. K.N analyzed RNAseq data under the supervision of S.M. H.T and K.P performed eVLP particle analysis and size characterization under the supervision of D.J.M. All authors read and approved the manuscript.

## MATERIAL AND METHODS

### Cell culture and cell lines

All cell lines were maintained at 37°C in a humidified incubator with 5% CO_2_ and routinely tested for mycoplasma contamination.

Lenti-X 293T (Takara 632180), HEK293T (ATCC CRL-1573), RPE-1 (ATCC CRL-4000) and A549 (ATCC CCL-185) cells were cultured in DMEM (Corning 10-013-CV) supplemented with 10% Fetalgro bovine growth serum (RMBIO FGR-BBT) and 1% penicillin/streptomycin (Cytiva Hyclone SV30010). HCT116 (ATCC CCL-247) and K562 (ATCC CCL-243) cells were maintained in RPMI (Corning 10-040-CV) supplemented with 10% Fetalgro bovine growth serum and 1% penicillin/streptomycin. HAP1 cells (Horizon Discovery C859) were cultured in IMDM (Gibco 12440053) supplemented with 10% Fetalgro bovine growth serum and 1% penicillin/streptomycin. Adherent cells were dissociated using 0.05% Trypsin-EDTA (Gibco 25300054).

Primary human T cells were isolated from healthy donors under ethics approval from the Health Research Ethics Board of Alberta (HREBA.CC-16-0762). Informed consent was obtained from all donors. T cells were purified using the EasySep Human T Cell Isolation Kit (STEMCELL Technologies 17951) and cultured in TexMACS medium (Miltenyi Biotec 130-097-196) supplemented with 5% Fetalgro bovine growth serum and 200 IU/mL recombinant human IL-2 (Miltenyi Biotec 130-097-743). T cells were activated with T Cell TransAct beads (Miltenyi Biotec 130-128-758) one day prior to transduction.

### Plasmid construction and cloning

All gRNAs and pegRNAs were synthesized as complementary oligonucleotides (IDT) and cloned into the appropriately digested plasmid vectors using the Golden Gate cloning, as previously described^88,8987,8888,89^. Briefly, oligonucleotides were phosphorylated using T4 polynucleotide kinase (NEB, M0201) and annealed by heating to 95°C for 5 minutes followed by slow cooling to room temperature. Successful cloning of (pe)gRNAs was verified by Sanger sequencing (Azenta Life Sciences).

Additional plasmids (described below) were constructed using Gibson Assembly and verified by whole plasmid sequencing (Azenta Life Sciences or Plasmidsaurus). The eVLP-TadCBEd construct (BB666) was generated by digesting pCMV-MMLVgag-3xNES-ABE8e (Addgene 181751) with BglII to remove Tad8 and nCas9 components. Mutated Tad8e^90^ and nCas9 were then reintroduced, the 2xUGIs were amplified from pCMV-ancBE4max (Addgene 112094). The PE2max-eVLP construct (BB499) was assembled by digesting pCMV-MMLVgag-3xNES-Cas9 (Addgene 181752) with BglII and inserting PE2max amplified from pCMV-PEmax (Addgene 174820). The Gag, NES sites, protease cleavage site, FLAG and SV40 NLS sequences were reconstituted by PCR amplification from the original plasmid. The plenti-PE2max-blastR construct (BB236) was created by digesting pLenti PE2-BSD (Addgene 161514) with EcoRI and AgeI, followed by insertion of PE2max amplified from pCMV-PEmax-P2A-BSD (Addgene 174821).

The pLenti MLH1dn dox-inducible plasmid was assembled by cloning Mlh1dn from pCMV-PEmax-P2A-hMLH1dn (Addgene 174828) into pRDA_174 (Addgene 136476) under an inducible promoter from XLone-GFP (Addgene 96930). A hygromycin resistance gene from mini pUC19 hygroR (BB224) was incorporated for selection. The Dox-inducible GFP eSTOP reporter (TAG) construct (BB299), and the gRNA ABE GFP-eSTOP (TAG) (BB298) were generous gifts from Dr. Ciccia (Columbia University). Complete oligonucleotide sequences are provided in Supplementary Tables S1 and S2, and additional published plasmids used in this study are listed in Supplementary Table S3.

### Lentivirus production and cell transduction

Lentiviruses were produced in Lenti-X 293T cells seeded at 5 x 10^5^ cells per well in 6-well plates with 2 mL complete DMEM. After 24 hours, cells were transfected with 3 µg total DNA comprising equal molar ratios of psPAX2 (Addgene 12260) and pMD2.G (Addgene 12259), along with the transfer vector. Transfection was performed 250 µL using OptiMEM (Gibco 31985070) and 3 µL of 1 mg/mL polyethylenimine (PEI) (Sigma 408727). The transfection mixture was incubated for 15 minutes at room temperature before being added dropwise to the cells. Following overnight incubation, media was replaced with 2 mL of fresh complete DMEM, and cells were incubated for an additional 48 to 60 hours. Viral supernatants were collected, centrifuged at 500 x g for 5 minutes, and filtered through 0.45 µm filters to remove cellular debris. Lentiviruses were used immediately or aliquoted and stored at −80°C. For large-scale screening applications, production was scaled up to 15 cm dishes using proportionally increased reagent volumes.

For lentiviral transduction, target cells were seeded at 3 x 10^5^ cells per well in 6-well plates with 2 mL of complete media. The following day, lentivirus was added to the cells along with 10 µg/mL polybrene. Cells were spinoculated at 935 x g for 2 hours at 32°C followed by a 12-hour incubation at 37°C and 5% CO_2_. Media was then replaced with fresh complete media, and antibiotic selection began on the next morning using puromycin (1 µg/mL), blasticidin (10 µg/mL), or hygromycin (100 to 400 µg/mL as determined by kill curves for each cell line).

### Production and characterization of eVLPs

eVLPs were generated in Lenti-X 293T producer cells following the established protocol^40^. Briefly, cells were seeded at 5 x 10^6^ cells per 75 cm² flask in 10 mL of complete DMEM, and transfections were performed 24 hours later using either JetPRIME (Polyplus 101000046) according to Banskota et al., 2022^40^, or PEI. For Targeting gRNA (T) loaded eVLPs, transfection was performed using JetPRIME with optimized ratios: 400 ng of VSV-g (BB14), 3.375 µg of pBS-CMV-gagpol (Addgene 35614), 1.125 µg of the editor plasmid, and 4.4 µg of the guide plasmid mixed with 18.6 µL of JetPRIME in JetPRIME buffer for a total volume of 500 µL^40^. The mixture was incubated for 10 minutes at room temperature before being added to the cells. Non-Targeting gRNA (NT) eVLPs were produced using PEI transfection with 860 ng of VSV-g, 7.258 µg of pBS-CMV-gagpol, 2.419 µg of the editor plasmid, and 9.462 µg of the (pe)gRNA plasmid (BB106, for gRNAs; or BB523 for pegRNAs). pegRNA-free (-) eVLPs were generated using PEI transfection with 1.634 µg of VSV-g, 13.791 µg of pBS-CMV-gagpol, and 4.597 µg of the editor plasmid. For PEI transfections, plasmid mixtures were combined with 60 µL of PEI (1 mg/mL) in OptiMEM for a total volume of 500 µL and incubated 15 minutes at room temperature before being added dropwise to the cells. Media was replaced with 10 mL of fresh complete DMEM at 12 hours post-transfection. eVLPs-containing supernatants were harvested 96 hours after transfection, centrifuged at 500 x g for 5 minutes and filtered. The eVLP supernatants were then concentrated 100-fold in serum-free, antibiotic-free media or PBS by ultracentrifugation at 30,000 x g for 2 hours at 4°C. Concentrated eVLPs were used immediately or stored at 4°C for up to one week. Particle concentration and size distribution were measured using TRPS on the IZON qNano TRPS Particle Analyzer (IZON Science Q5013) according to manufacturer’s instructions. eVLP samples were initially diluted 1:50 in PBS, then further diluted (1:10 to 1:1,000) in PBS containing 0.1% TWEEN-20 (Sigma-Aldrich P1379) for quantification. Each sample were analyzed alongside IZON calibration beads (IZON Science CPC100) using a 50-330 nm Nanopore (IZON Science NP100) until 1,000 events were recorded. Data were processed using the IZON Control Suite software (v3.2) to calculate particle concentration and size distribution.

### Base editing and prime editing via transfection

Editing experiments were performed in HEK293T cells using two distinct transfection protocols. For *Trans*IT-293 (mirusbio MIR 2700) transfections, cells were seeded at 70-80% confluency in a 48-well plates at 5.5 x 10^5^ cells per well in 250 µL complete DMEM according to the manufacturer’s instructions^88,91^. For Lipofectamine 2000 (Invitrogen 11668027) transfections, cells were seeded in 96-well plate with 1.8 x 10^5^ per well in 100 µL complete DMEM as previously described^50^.

Transfections were performed 24 hours after cell seeding. For base editing experiments, cells were transfected with 387 ng of TadCBEd plasmid (Addgene 193835) or ABE8e plasmid (Addgene 138489) and 113 ng of gRNA plasmid using 1 µL of *Trans*IT-293 reagent. For prime editing, cells were transfected with 417 ng of PEmax plasmid (Addgene 174828) and 83 ng of pegRNA plasmid. *Trans*IT-293 DNA transfection mixtures were incubated for 20 minutes at room temperature before dropwise addition to cells. Lipofectamine 2000 transfections consisted of 292 ng of base or prime editor plasmids with 96 ng of the (pe)gRNA plasmid and 1 µL of Lipofectamine 2000 reagent. The DNA:lipid complexes were incubated for 10 minutes at room temperature before being added dropwise to the cells. Cells were maintained in culture for 72 hours until collection by centrifugation at 300 x g for 3 minutes. Cell pellets were stored at −20 °C until genomic DNA extraction.

### Base editing and prime editing via eVLP transduction

For eVLP-mediated editing, HEK293T cells were seeded in 48-well plates at 3.5 x 10^5^ cells per well in 250 µL of complete DMEM. Cells were transduced 8 hours after seeding with 100 µL of concentrated eVLPs. Base editing experiments used ABE8e or TadCBEd eVLPs loaded with appropriate gRNA, while prime editing utilized PE2max eVLPs loaded with the relevant pegRNAs. Cells were collected 72 hours post-transduction, and cell pellets were stored at −20°C until genomic DNA extraction.

### Multiple eVLP transductions

For compound transduction experiments, HEK293T, HAP1-MLH1dn, RPE-1-MLH1dn or K-562-MLH1dn cells were seeded in a 48-well plate at 3.5 x 10^5^ cells per well with 250 µL of complete media^40^. Cells were transduced 8 hours later with either 25 or 100 µL of PE2max-eVLP loaded with specific pegRNAs including *PCNA* +2 AA>GC (BB146); HEK3 +1 CTTins (BB415); HEK3 + 1T del (BB416); *RNF2* +5 G>T (BB732); *ADA* +9 C>T (BB854) or *EMX1* + 1 G>T (BB926). eVLPs were added directly to the culture media.

For time-course analysis, cells were collected 1, 2, 3, 4 or 5 days after transduction and were stored at −20°C until genomic DNA extraction. For multiple transduction experiments, cells were trypsinized, counted and replated at 3.5 x 10^5^ cells per well (48-well plates) 8 hours before subsequent addition of eVLPs. To investigate the days between transduction, cells that underwent four sequential transductions were collected at the indicated times (days 4, 8, 12, 16, or 20 post-initial transduction).

### PRIME-VLP protocol

The optimized PRIME-VLP protocol for HEK293T involves transducing 3.5 x 10^5^ cells with 25 µL (for targeted editing) to 50 µL (for screen) of eVLP (700 to 1,400 cells/µL), every three days. Cells are collected 3 days after the final transduction, on day 12 or 24 for the screens. eVLP volumes were adjusted according to cell line-specific optimization and scaled proportionally to the number of cells seeded in each experiment.

### Genomic DNA extraction, Sanger and NGS

Genomic DNA was extracted from the frozen cell pellets using QuickExtract DNA Extraction Solution (Biosearch Technologies QE090050) as described previously^91^. Editing in samples was assessed using either Sanger or NGS (Azenta Life Sciences). For Sanger sequencing analysis, samples were amplified using locus-specific primers in the Supplementary Table S4. Chromatogram files (.ab1) were used to quantify editing efficiency using EditR^92^ v1.1.10.

For NGS analysis, samples underwent dual PCR amplification. The first PCR used Q5 High-Fidelity DNA Polymerase (NEB M0491L) and locus-specific oligonucleotides listed in the Supplementary Table S5. A second PCR was performed using 1 µL of the first PCR as template to incorporate Illumina indexes and adapter sequences. Amplicon libraries were sequenced on an Illumina NovaSeq platform using paired-end 2 x 150 bp reads with a 5% PhiX spike-in. Sample demultiplexing and FASTQ generation were performed using Illumina’s bcl2fastq tool with standard indexing primers. Editing at endogenous loci was evaluated using CRISPResso2 (v2.3.2)^93^, with a Phred quality cut-off of ≥20. Reads were aligned to reference amplicons for quantitative analysis. Prime editing efficiency was quantified within an 11-bp window centred on the Cas9 (H840A) nick site, using the pegRNA spacer and 3’-extension sequences. Base editing efficiency was assessed using quantification_window_center of −7 and a quantification_window_size of 20; the resulting Alleles_frequency_table.txt was then used to determine editing efficiency.

The accession numbers for the NGS data are SRA: SUB15455217; BioProject PRJNA1294868.

### Cell viability analysis

Each sample was counted in triplicate using the DeNovix CellDrop counter to determine viability. HEK293T were seeded as described previously, while T cells were plated in round-bottom 96-well plates at 5 x 10^5^ per well in 50 µL complete TexMACS media. Cells were either left untreated or transduced 8 hours later with 25 µL (PRIME-VLP protocol) or 100 µL of eVLPs. Both HEK293T and T cells were counted in triplicate and replated at initial cell densities on days 3, 6 and 9. PRIME-VLP cells received four sequential transductions of 25 µL eVLPs each, totalling 100 µL eVLPs from a single preparation. Final cell counts were performed on day 12 for all experimental conditions.

### Evaluation of sequential eVLP transduction efficiency

HEK293T ABE reporter cells containing a doxycycline-inducible GFP cassette with a premature stop codon were seeded in a 48-well plate at 3.5 x 10^5^ cells per well with 250 µL of complete DMEM. Eight hours post-seeding, cells were transduced with 25 µL of ABE8e eVLPs carrying gRNA targeting the stop codon. Three days after plating, 1 µg/mL of doxycycline was added. After 24 hours, cells were sorted for GFP expression to obtain a homogeneous population of GFP-positive cells (Cumming School of Medicine Flow Cytometry Core Facility). Immediately after sorting, GFP-positive cells were replated in 48-well plates at 3.5 x 10^5^ cells per well. Non-edited, GFP-negative HEK293T reporter cells were plated in parallel as secondary transduction controls. When cells were fully attached (1 hour), both populations were transduced with 25 µL of TadCBEd-eVLPs carrying RASA2-targeting gRNA. Cells were collected 72 hours post-transduction, gDNA was extracted, and *RASA2* editing efficiency was quantified by Sanger sequencing.

### Off-target editing analysis

Off-target editing analysis was performed using HEK293T cells transduced with eVLP either containing Cas9 or PE2max loaded with HEK3- and HEK4 (pe)gRNAs, following the standard eVLP or PRIME-VLP transduction protocols detailed above. For comparison with conventional transfection, HEK293T cells were seeded in 48-well plates at 5.5 x 10^5^ cells per well in 250 µL complete DMEM for transfection using *Trans*IT-293 reagent, according to the manufacturer’s protocol. Briefly, transfection mixtures contained 750 ng of Cas9 (Addgene 42230) and 250 ng of HEK3 (BB867) or HEK4 (BB868) gRNA plasmids mixed with 2 µL of *Trans*IT-293 in 25 µL total volume adjusted with OptiMEM. The DNA:lipids complexes were incubated for 10 minutes at room temperature before being added to HEK293T cells. All cells were collected on day 12 and stored at −20 °C until gDNA extraction. On- and off-target editing frequencies at HEK3 and HEK4 loci were quantified by NGS.

### RNA sequencing and transcriptome analysis

Primary human T cells were isolated from four healthy donors and seeded in round-bottom 96-well plates at a density of 5.0 x 10^5^ cells per well in 50 µL of complete TexMACS medium. Cells were either left untreated as controls or subjected to transduction using either the eVLP or PRIME-VLP protocols, as detailed above. Following treatment, T cells were harvested on day 3 for assessment of acute cellular responses and day 12 for long-term responses. All samples were immediately frozen at −80°C prior to RNA extraction and sequencing, both performed by Azenta Life Sciences. Twenty-four RNA-Seq libraries were generated with high-quality metrics, averaging ∼37 million reads per library (range: 25-50 million reads) using 150-nucleotides paired-end sequencing. Data quality assessment was performed using FastQC, with all samples passing QC with a mean base quality score of >35. Raw FASTQ files were aligned to the GRCh38 human reference genome using the STAR aligner^94^ in two-pass mode, with “–quantMode” parameter configured to generate read counts per gene. The alignment yielded > 80% uniquely mapped reads across all samples. Gene annotations were performed using the gencode.v44chr_patch_hap1_scaff.annotation.gtf file, with resulting gene counts were utilized for subsequent differential expression analysis. Differential gene expression analysis was conducted using DESeq2^95^. Comparisons were made between treated samples (1 x 100 µL or PRIME-VLP) and untreated controls, analyzed independently for day 3 and day 12 time points. Genes were considered differentially expressed when meeting criteria of BH-adjusted p-value < 0.05 and absolute |Log2 fold change| >1. Pathway enrichment was performed using fGSEA^57^ with previously reported CAR T-cell gene sets that represent distinct aspects of CAR-T cell biology^96^.

### Construction and cloning of the miniature sensor pegRNA library

The miniature sensor pegRNA library consisted of 31 pegRNAs, each with manually designed target sequences. These constructs were cloned into a modified lentiGuide-Puro (BB26), that had been pre-modified to remove the original gRNA scaffold sequence and incorporate the epegRNA 3’ extension sequence along with two restriction enzyme sites. Individual pegRNAs were inserted into BsmBI-linearized BB770 using Gibson assembly. Target sequences were subsequently cloned by sequential digestion with EcoRI and BamHI. All constructs were verified by Sanger sequencing (Azenta Life Sciences).

Sequences of the pegRNAs and targets are listed in the supplementary Tables S2 and S6.

### Pooled prime editing mini screen and analysis

Cell lines were infected with the miniature sensor pegRNA library using lentivirus at an MOI = 0.3 with a representation of 1,000x (>3.1 x 10^5^). Cells were selected with 1 µg/mL puromycin for a minimum of 3 days. After recovery, 3.5 x 10^5^ cells were harvested and frozen as an initial screen sample, while an equivalent number of cells were divided into experimental groups. One group received PE2max (BB236) lentivirus at a MOI of 1 followed by selection with 10 µg/mL blasticidin for at least 5 days. A second group was transduced with either Non-Targeting pegRNA (NT) (BB523) loaded PE2max-eVLP or pegRNA-free (-) PE2max-eVLPs following the PRIME-VLP protocol. The PRIME-VLP transduction protocol involved seeding cells in a 48-well plate at 3.5 x 10^5^ cells per well with 250 µL complete medium, followed by eVLP transduction 8 hours post-seeding using optimized eVLP amounts. Cells were subsequently trypsinized, counted and re-transduced every 3 days until day 24, completing a total of 8 transduction cycles. Lentivirus-infected control cells were passaged in parallel with PRIME-VLP treated cells throughout the 24-day experimental period. All MLH1dn cell lines were maintained in 1 µg/mL doxycycline starting from PE2max infection and first PRIME-VLP transduction through the duration of the screens. Cell pellets containing a minimum of 3.5 x 10^5^ cells per condition were collected on days 6, 15 and 24 (final) and stored at −20°C until genomic DNA extraction.

Genomic DNA extraction was performed using standard protocols, followed by a 2-step PCR amplification strategy. First, pegRNAs and targets were amplified using NEBNext Ultra II Q5 Master Mix (NEB M0544X). A second PCR reaction was performed using the first PCR as a template, adding adapters and barcodes sequences, as described previously^97^. Oligonucleotides used for the PCR amplification are available in the supplementary Table S5. Amplicon libraries were sequenced by Azenta Life Sciences using the Illumina NovaSeq platform with paired-end 2 x 150 bp reads with a 5% PhiX spike-in. Sample demultiplexing and FASTQ file generation were performed using Illumina’s bcl2fastq tool with standard indexing primers. Analysis of miniature sensor library data was conducted using a custom R script that searched sequence reads for target-specific sensor sequences, categorized reads by target, and calculated editing rates by quantifying reads that matched either the wild-type or intended edit sequence. Targets demonstrating < 1% editing across all the experimental conditions were excluded from downstream analysis.

### PE2max expression and RT-qPCR

Multiple cell lines including HAP1-MLH1dn, HCT116, K562-MLH1dn, A549-MLH1dn, and HEK293T were infected in duplicate with PE2max lentivirus at approximately MOI of 1. Following blasticidin selection at 10 µg/mL, initial cell pellets were collected and frozen at −80°C as baseline samples for RNA extraction. Cells were maintained in continuous culture with regular passaging for 24 days, after which final samples were collected for comparative RNA analysis.

RNA extraction was performed by TRIzol chloroform extraction. Briefly, cell pellets were resuspended in TRIzol at a volume of 1 mL per 1.0 x 10^7^ cells, incubated for 5 minutes at room temperature, and centrifuged at 13,000 x g for 5 minutes at room temperature. Chloroform was added at one-fifth the initial TRIzol volume (e.g., 200 µL per mL TRIzol). Samples were vortexed for 15 seconds, incubated for 3 minutes at room temperature, and centrifuged at 12,000 x g for 15 min at 4°C. The aqueous phase was carefully collected, and RNA was precipitated by adding 100% isopropanol at half the original volume (e.g., 500 µL per mL TRIzol) followed by incubation at room temperature for 10 min and centrifugation at 7,500 x g for 5 min at 4°C. Pellet was air dried and resuspended in DEPC-treated water. RNA quantification was performed using NanoDrop spectrophotometer.

Reverse transcription was performed with the SuperScript IV First-Strand Synthesis System (Invitrogen 18091050) with random hexamer primers according to manufacturer specifications. Reactions were performed at room temperature for 10 minutes, 50°C for 1 hour, and 80°C for 10 minutes. The resulting cDNA was diluted 1:20 for reactions using 1 µg RNA input or 1:4 for lower RNA input. Quantitative PCR was conducted using Applied Biosystems QuantStudio 6 with reactions containing 5 µL of 2X *Power* SYBR Green PCR Master Mix (Applied Biosystems 4367659), 0.1 µL each primer (100 µM) (primer sequences are available in the supplementary Table S7), and 1 µL diluted cDNA in 10 µL total volume. All samples were analyzed in triplicate. The qPCR program was as follows: Initial hold stage at 50°C for 2 minutes and 95°C for 10 minutes, followed by 40 cycles of 95°C for 15 seconds and 60°C for 30 seconds, with a final melt curve analysis at 60°C for 1 minute ramping to 95°C for 5 seconds.

### Prime editor silencing and editing

HEK293T cells were infected in duplicate with PE2max (BB236) lentivirus at an MOI of 1 and selected with 10 µg/mL blasticidin. One set of cells was immediately frozen while another was maintained in continuous culture for 24 days. Low-passage PE2max HEK293T cells were thawed on day 21, and both low and high-passage cells were transfected on day 24 with pegRNAs (BB520, BB676, and BB677). After 72-hour incubation, all cells were pelleted and stored at −20°C prior to genomic DNA extraction. Editing efficiency was assessed through locus-specific PCR amplification followed by Sanger sequencing (Azenta Life Sciences).

### Construction and cloning of p53/Nutlin library

The p53 Nutlin sensor library was adapted from Gould *et al*., 2024^29^ and synthesized by Twist Biosciences as a lyophilized oligonucleotide pool. Library amplification and cloning were performed according to established protocols, using the lyophilized material resuspended in TE buffer (pH 8.0) at 10 ng/µL and subsequently diluted to 1 ng/µL. Ten parallel PCR reactions were performed using 1 ng of oligonucleotide pool per reaction for 14 cycles total (primers available in Supplementary Table S1). All PCR reactions were pooled and purified using AMPure XP beads (Beckman Coulter A63881).

The sensor library backbone (BB1131) was derived from lentiGuide-Puro (Addgene 52963) with modifications that included the removal of the gRNA scaffold sequence and the addition of a BsmBI site (oligonucleotides are available in Supplementary Table S1). Both the amplified library and backbone were digested with BsmBI (NEB R0734S) and EcoRI-HF (NEB R3101S) and gel purified using GenepHlow Gel/PCR kit (Geneaid DFH300) and the AMPure XP beads. Ligation was performed using the High concentration T4 DNA ligase (NEB M0202M) at 25°C overnight, with 7:1 insert-to-vector ratio. Following heat inactivation at 65°C for 10 minutes, four electroporation reactions were conducted using 25 µL BioSearch Technologies Endura DUOs Electrocompetent cells (60242-2) with 3 µL ligation product per reaction (Bio-Rad MicroPulser electroporator 1652100). Bacteria cultures were grown overnight at 32°C in 2 L of LB medium with 100 µg/mL of carbenicillin.

Library representation was verified by Amplicon-EZ sequencing, and recombination rates were confirmed by Sanger sequencing of 10 individual colonies.

### p53/Nutlin library screening and analysis

Both HEK293T and A549-MLH1dn screens were performed in duplicate following established protocols^29,97^. Briefly, initial library infection involved 1 x 10^7^ cells transduced with *TP53* sensor library lentivirus at MOI 0.3, followed by puromycin selection at 1 µg/mL. Five to six days post-transduction, 3.5 x 10^6^ selected cells, representing > 500x library coverage, were either collected as initial samples or plated for experimental conditions: no-editor controls plated in 15 cm dishes and PE2max lentivirus infection, and NT pegRNA-loaded PE2max-eVLP transduction conditions seeded at 2.5 x 10^5^ cells per well in 6-well plates, for a total of 14 wells each.

Lentivirus infections were performed 24 hours post-seeding with plenti-PEmax-blastR (BB236) at MOI 1; followed by blasticidin selection at 10 µg/mL for at least 5 days. A549-MLH1dn cells were maintained in 1 µg/mL doxycycline throughout the experiment beginning on day 0. Every three days for 24 days, 3.5 x 10^6^ cells from each condition were replated, with eVLP-transduced cells receiving additional transductions for a total of eight transductions over 24 days. Cell pellets of 3.5 x 10^6^ cells were collected and stored at −20°C for each condition and time point.

The HEK293T screen concluded on day 24 with final sample collection of 3.5 x 10^6^ cells. On day 24, 3.5 x 10^6^ A549-MLH1dn cells were collected for each condition and replated for two additional days of expansion before being separated into untreated and Nutlin-3 treatment groups. Nutlin-3 (Selleckchem S1061) was added to the cells on day 27 at 10 µM and maintained for 11 days until screen completion of the screen on day 38. Cells were passaged every 3 days, maintaining >500X representation. Cell pellets were collected and frozen at −20°C for day 5 and day 11 (final) of Nutlin-3 treatment. Genomic DNA extraction was performed using the Qiagen DNeasy Blood & Tissue kit (Qiagen 69504) according to the manufacturer’s protocols. NGS samples were prepared using dual PCR amplification with NEBNext Ultra II Q5 Master Mix (NEB M0544X). Initial PCR amplification consisted of 19 cycles using total genomic DNA from each sample as a template across 14 reactions with 3 µg of template genomic DNA. The second PCR of 11-16 cycles added adapters and sample barcodes using pooled PCR1 products across 4 reactions per sample (primers listed in Supplementary Table S5). Final PCR products were pooled, purified, and sequenced using NovaSeq with paired-end 2 x 150 bp reads and 5% PhiX spike-in.

Sample demultiplexing and FASTQ file generation were performed using Illumina’s bcl2fastq tool with standing indexing primers. Editing frequency analysis employed a custom R script that combined forward and reverse reads, aligned them to specific pegRNAs using 3’ extensions, and calculated editing percentages using the formula (Edited Reads / (Edited Reads + Wild-type Reads)) x 100. Functional screen analysis utilized the MAGeCK (v0.5.9.5) pipeline^98^ to estimate statistical significance and fold change for each pegRNA using default parameters. Statistical comparisons assessed differences between Nutlin-3-treated and control groups on day 11, with results merged with editing efficiency based on pegRNA identifiers.

## Notes

### Competing Interest Statement

The authors have declared no competing interest.

