## Supplementary for "Compound Delivery of eVLPs Enhances Prime Editing for Targeted Genome Engineering and High-Throughput Screening"

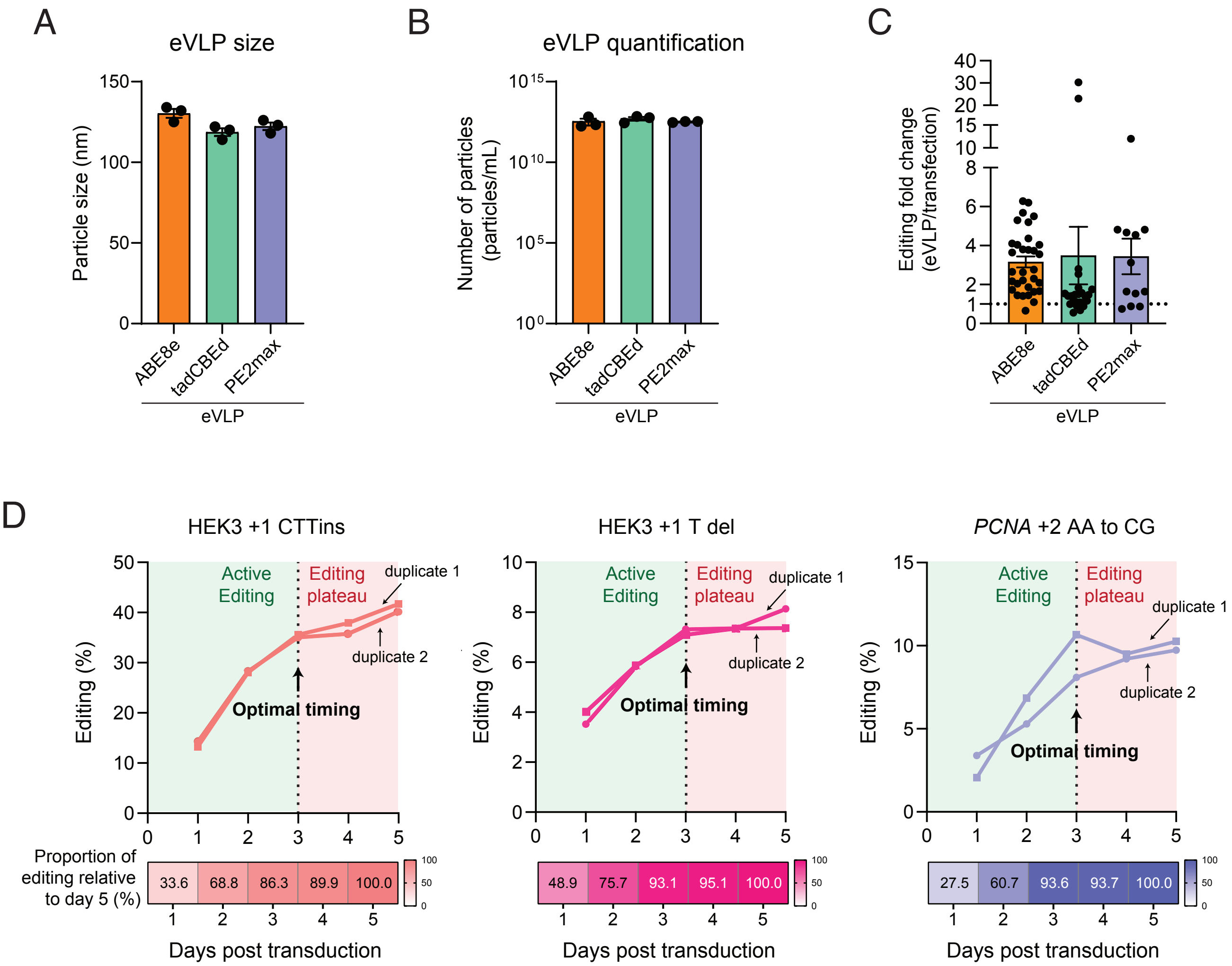

A
eVLP size
Particle size (nm)
150
100
50
0
ABE8e
tadCBEd
PE2max
eVLP
B
eVLP quantification
Number of particles (particles/mL)
$10^{15}$
$10^{10}$
$10^{5}$
$10^{0}$
ABE8e
tadCBEd
PE2max
eVLP
C
Editing fold change (eVLP/transfection)
40
30
20
15
10
8
6
4
2
1
0
ABE8e
tadCBEd
PE2max
eVLP
D
HEK3 +1 CTTins
HEK3 +1 T del
*PCNA* +2 AA to CG
Active Editing
Editing plateau
duplicate 1
duplicate 2
Optimal timing
Editing (%)
Proportion of editing relative to day 5 (%)
33.6
68.8
86.3
89.9
100.0
48.9
75.7
93.1
95.1
100.0
27.5
60.7
93.6
93.7
100.0
Days post transduction


Figure S1

**Figure S1. eVLP characterization and editing kinetics**

**(A)** eVLP production consistency across editor types. Modal particle size measurements determined by TRPS demonstrating uniform size distribution between 100-150 nm across all editor types. Data represent individual preparations with mean ± s.e.m. from n = 3 independent productions.

**(B)** Particle quantification by TRPS showing comparable concentrations ($\sim 10^{12}$ particles/mL) for ABE8e, TadCBEd, and PE2max eVLPs.

**(C)** Fold-change analysis of eVLP versus transfection efficiency across all genome editors. Each point represents an independent edit (target and edited base, from Figure 1A-C) comparing eVLP delivery to transfection methods. Horizontal dotted line at y = 1 represents equivalent performance; all points above indicate eVLP superiority. Data show consistent 2-8-fold improvements for eVLP delivery across ABE8e, TadCBEd, and PE2max editors.

**(D)** Detailed kinetic analysis of prime editing for individual genomic targets. Line graphs show editing progression over 5 days for three targets (HEK3 +1 CTTins, HEK3 +1 T del, PCNA +2 AA to CG), with heat maps below displaying the cumulative proportion of editing relative to day 5 (color scale: white = 0%, dark = 100%). Green shading highlights the active editing window (days 1-3) where most total editing occurs. Vertical dashed lines indicate optimal timing for subsequent transductions. Data represent mean from n = 2 independent experiments.

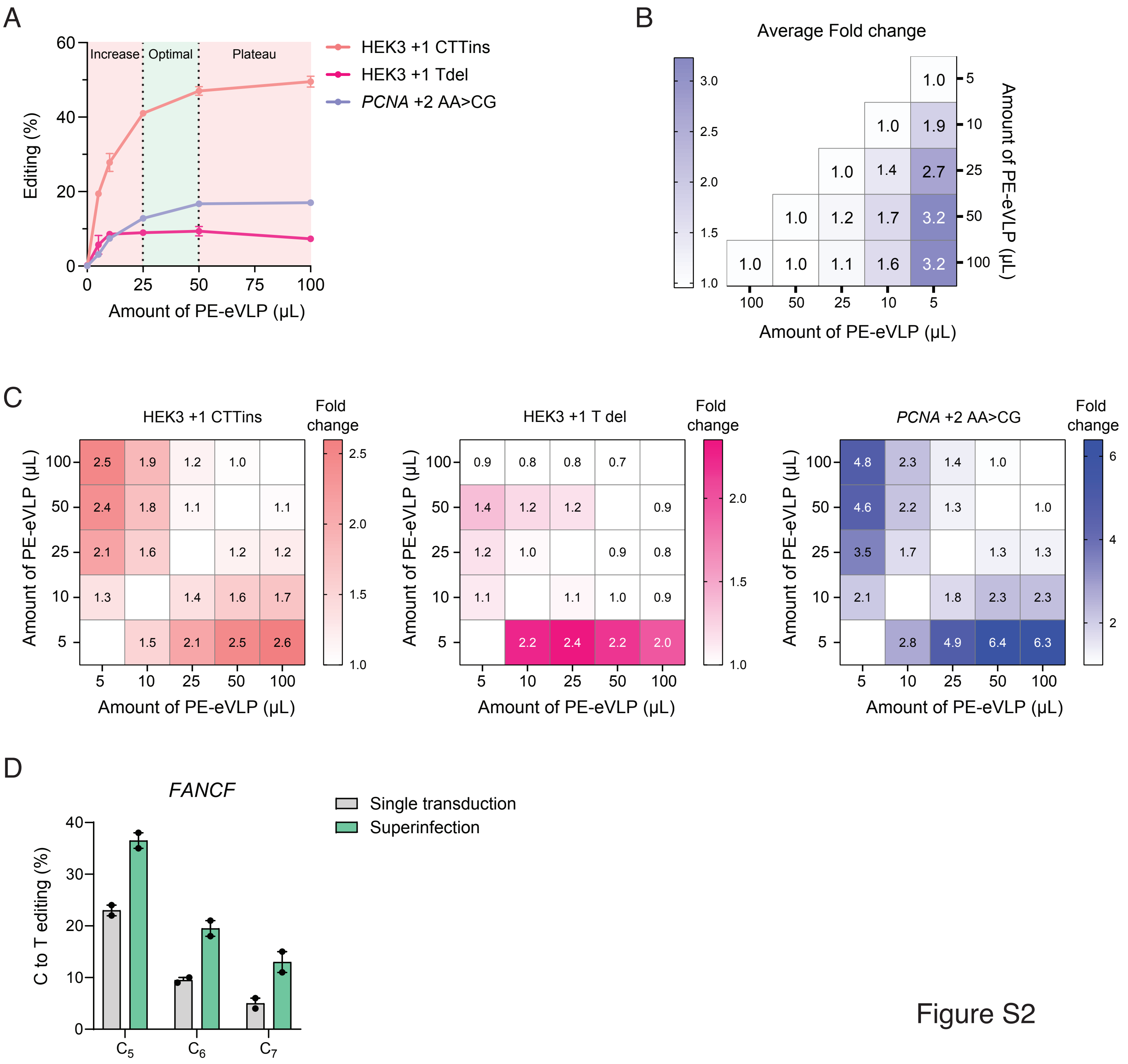
A
Increase
Optimal
Plateau
Editing (%)
Amount of PE-eVLP (μL)
HEK3 +1 CTTins
HEK3 +1 Tdel
PCNA +2 AA>CG
B
Average Fold change
Amount of PE-eVLP (μL)
Amount of PE-eVLP (μL)
C
HEK3 +1 CTTins
Fold change
HEK3 +1 T del
Fold change
PCNA +2 AA>CG
Fold change
Amount of PE-eVLP (μL)
Amount of PE-eVLP (μL)
D
FANCF
C to T editing (%)
Single transduction
Superinfection
C5
C6
C7

Figure S2

**Figure S2. Dose-response optimization and superinfection validation**

**(A)** Dose-response analysis across three prime editing targets (HEK3 +1 CTTins, HEK3 +1 T del, *PCNA* +2 AA>CG). Line graphs demonstrate non-linear dose-response relationships with distinct phases: suboptimal editing at low doses (pink shading, left), optimal editing efficiency in the 25-50 μL range (green shading, center), and saturation plateau at high doses (pink shading, right). Vertical dashed lines delineate the transition between increase, optimal, and plateau phases.

**(B)** Quantitative analysis of dose-response relationships averaged across all three targets. Heat map displays fold-change values calculated as editing efficiency at y-axis dose divided by editing efficiency at x-axis dose. Values >1.0 (blue) indicate improved editing with higher doses, while values approaching 1.0 (white) indicate saturation. The gradient demonstrates diminishing returns above 25 μL, supporting the use of sub-saturating doses for PRIME-VLP.

**(C)** Target-specific dose-response relationship for individual genomic loci. Heat maps show fold-change relationships for HEK3 +1 CTTins (left), HEK3 +1 T del (center), and *PCNA* +2 AA>CG (right). Each matrix compares editing efficiency between different dose combinations, with numerical values indicating fold-change improvements. Color intensity reflects the magnitude of improvement, with saturation effects evident at higher doses across all targets.

**(D)** Validation of sequential eVLP transduction in primary human T cells. Bar graph compares single transduction (grey) versus sequential transduction/superinfection (green) using TadCBEd-eVLPs targeting three independent loci (FANCF C5, C6, C7). Sequential dosing achieves superior editing efficiency, demonstrating that primary cells with intact innate immune responses remain permissive to multiple eVLP deliveries. Data represent mean ± s.e.m. from n = 2 independent experiments.

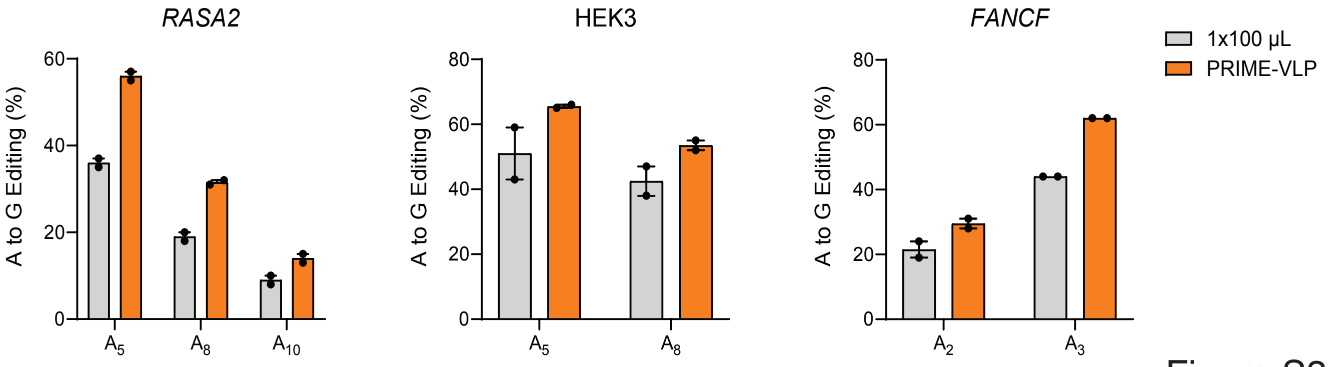

RASA2
HEK3
FANCF
A to G Editing (%)
0
20
40
60
80
A5
A8
A10
A2
A3
1x100 µL
PRIME-VLP


Figure S3

**Figure S3. PRIME-VLP improves base editing efficiency in primary human T cells**
Base editing efficiency comparison between single high-dose (1x100 µL, grey) and PRIME-VLP (4x25 µL, orange) delivery using ABE8e-eVLPs across three endogenous genomic loci (*RASA2*, HEK3, *FANCF*) in primary human T cells. PRIME-VLP consistently achieves superior editing efficiency across all targets and guide RNAs tested, with improvements ranging from 1.2 to 1.7-fold. The consistent enhancement demonstrates that PRIME-VLP benefits extend beyond prime editing to other CRISPR-based platforms and function effectively in primary cells with intact DNA repair mechanisms. Data represent mean ± s.e.m. from n = 2 independent experiments using T cells from different donors.

A

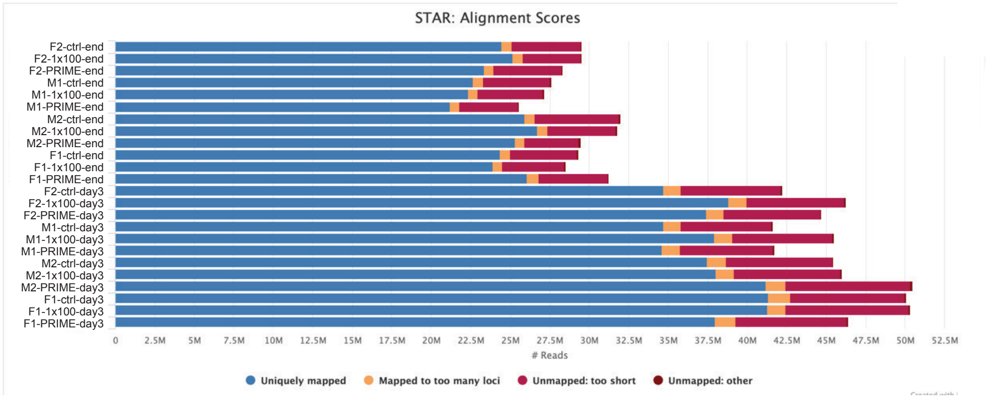


B

| CAR-T cell panel fGSEA: 1 x 100 µL | | | | | | | |
|---|---|---|---|---|---|---|---|
| **pathway** | **pval** | **padj** | **log2err** | **ES** | **NES** | **size** | **leadingEdge** |
| Activation | 0.97003 | 0.97003 | 0.00801916 | 0.140331 | 0.216161 | 2 | FOSB,FOS |
| Apoptosis | 0.7812188 | 0.97003 | 0.02414479 | 0.25 | 0.4946414 | 1 | FOS |
| Exhaustion | 0.7812188 | 0.97003 | 0.02414479 | 0.25 | 0.4946414 | 1 | FOS |
| Innate-like T-cells | 0.7812188 | 0.97003 | 0.02414479 | 0.25 | 0.4946414 | 1 | FOS |
| MAPK and PI3K Signaling | 0.7812188 | 0.97003 | 0.02414479 | 0.25 | 0.4946414 | 1 | FOS |
| Phenotype | 0.7812188 | 0.97003 | 0.02414479 | 0.25 | 0.4946414 | 1 | FOS |
| TCR signaling | 0.97003 | 0.97003 | 0.00801916 | 0.140331 | 0.216161 | 2 | FOSB,FOS |
| Toxicity | 0.6143856 | 0.97003 | 0.03614919 | 0.3333333 | 0.6595218 | 1 | EGR1 |
| Type1 interferon signaling | 0.6143856 | 0.97003 | 0.03614919 | 0.3333333 | 0.6595218 | 1 | EGR1 |
| **CAR-T cell panel fGSEA: PRIME-VLP** | | | | | | | |
| **pathway** | **pval** | **padj** | **log2err** | **ES** | **NES** | **size** | **leadingEdge** |
| Activation | 0.8971029 | 0.8971029 | 0.01545138 | 0.2 | 0.3435039 | 2 | FOS,FOSB |
| Apoptosis | 0.8461538 | 0.8971029 | 0.01945417 | 0.1666667 | 0.3284072 | 1 | FOS |
| Exhaustion | 0.8461538 | 0.8971029 | 0.01945417 | 0.1666667 | 0.3284072 | 1 | FOS |
| Innate-like T-cells | 0.8461538 | 0.8971029 | 0.01945417 | 0.1666667 | 0.3284072 | 1 | FOS |
| MAPK andPI3K Signaling | 0.8461538 | 0.8971029 | 0.01945417 | 0.1666667 | 0.3284072 | 1 | FOS |
| Phenotype | 0.8461538 | 0.8971029 | 0.01945417 | 0.1666667 | 0.3284072 | 1 | FOS |
| TCR signaling | 0.8971029 | 0.8971029 | 0.01545138 | 0.2 | 0.3435039 | 2 | FOS,FOSB |

C

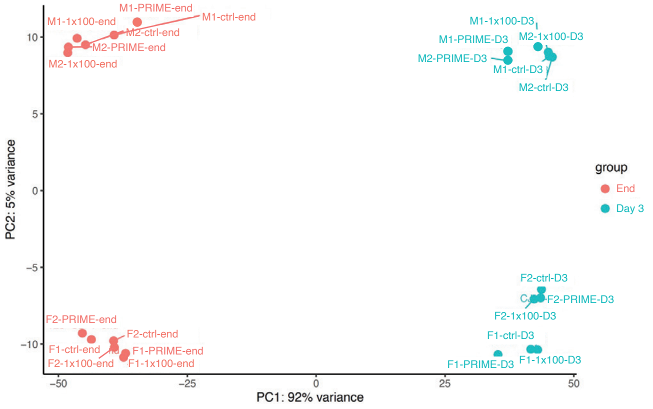


D

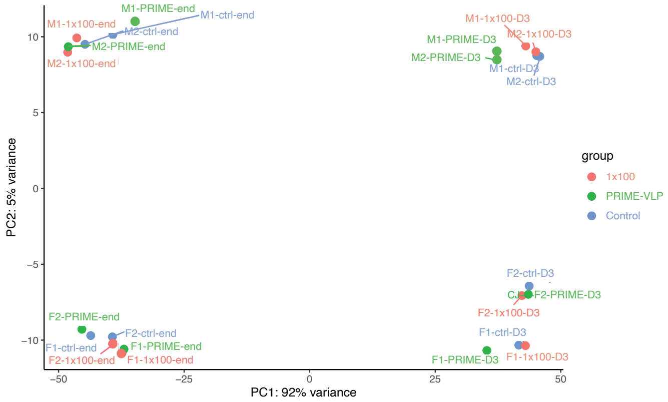


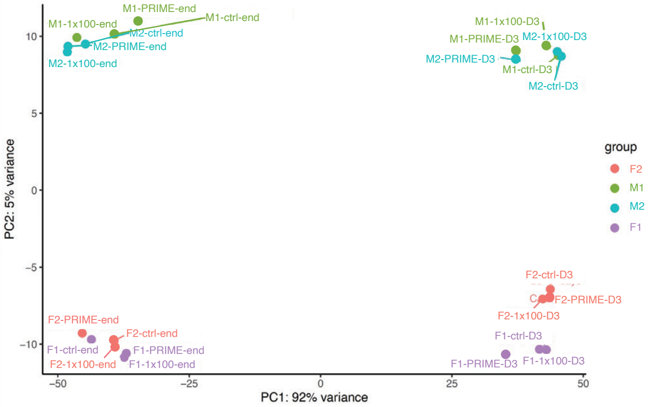


Figure S4

**Figure S4. Comprehensive transcriptomic analysis**

**(A)** RNA sequencing quality metrics and alignment statistics. STAR alignment summary from RNA-seq libraries generated from 4 healthy donors (M1 and M2 = male; F1 and F2 = female) across three conditions (ctrl: untreated control, 1x100: single high-dose, PRIME: PRIME-VLP) and two timepoints (day 3 and day 12). All samples achieved >80% unique mapping efficiency with an average sequencing depth of 37 million reads per library and high-quality metrics (mean base quality >35), confirming robust data quality.

**(B)** Gene Set Enrichment Analysis (fGSEA) results for CAR-T cell-specific pathways. Tables compare pathway enrichment between single high-dose (1x100 μL, top) and PRIME-VLP (bottom) conditions relative to untreated controls. Statistical parameters include p-values, adjusted p-values (padj), log2 error rates, enrichment scores (ES), normalized enrichment scores (NES), pathway sizes, and leading edge genes. No pathways showed significant enrichment (padj < 0.05) in either condition, indicating minimal pathway-level perturbations from eVLP treatments.

**(C-D)** Principal component analysis (PCA) demonstrating the relative contributions of experimental variables to transcriptional variance. Each point represents one biological sample, with colors indicating treatment groups (red: untreated control, blue: PRIME-VLP, green: single high-dose). (C) Day 3 timepoint and (D) day 12 timepoint analyses show that samples cluster primarily by donor and timepoint rather than treatment method. The variance explained by PC1 (92%) and PC2 (5%) indicates that temporal dynamics and inter-donor variability have substantially greater impact on gene expression than delivery method, highlighting the minimal transcriptional perturbation caused by eVLP treatments.

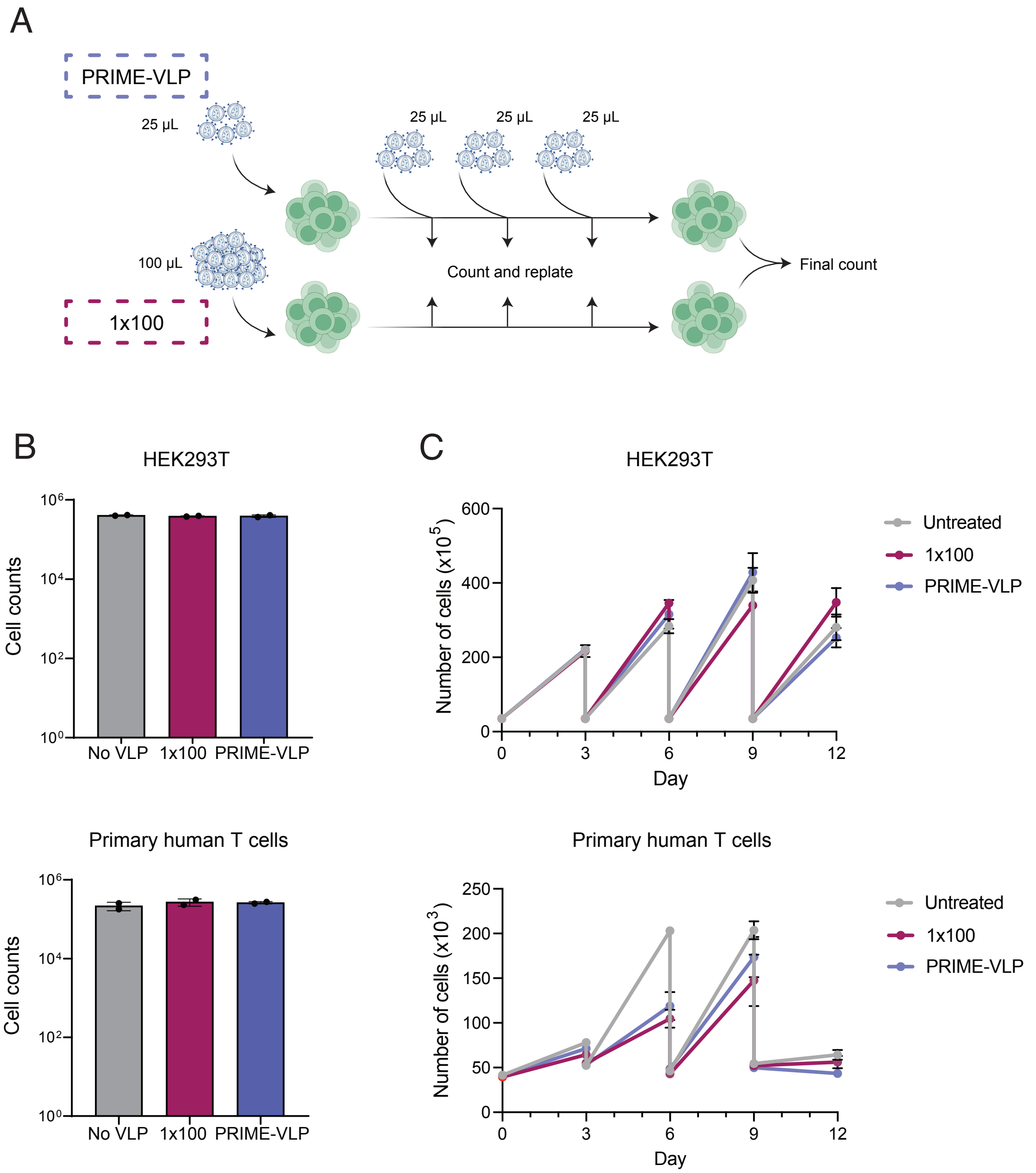
A
PRIME-VLP
25 µL
25 µL
25 µL
25 µL
100 µL
1x100
Count and replate
Final count
B
HEK293T
Cell counts
No VLP
1x100
PRIME-VLP
Primary human T cells
C
Number of cells (x10^5)
Day
Untreated
1x100
PRIME-VLP
Number of cells (x10^3)

Figure S5

**Figure S5. Cell viability and proliferation analysis**

**(A)** Experimental workflow schematic for viability assessment. Both HEK293T cells and primary human T cells were subjected to three treatment conditions: untreated controls (grey), single high-dose (1x100 μL, red), or PRIME-VLP (4x25 μL, blue). Cells were counted using automated cell counting and replated at standardized densities every 3 days throughout the 12-day experimental period to assess both acute and sustained viability effects during the complete PRIME-VLP treatment course.

**(B)** Cell viability quantification at day 12 for HEK293T cells (top) and primary human T cells (bottom). Bar graphs show no significant differences in final cell counts between untreated controls and either eVLP treatment regimen, demonstrating that neither PRIME-VLP nor single high-dose delivery adversely affects cell survival. Data represent mean ± s.e.m. from n = 2 independent experiments.

**(C)** Longitudinal cell proliferation tracking over the 12-day treatment period. Line graphs show cell expansion kinetics for HEK293T cells (top) and primary human T cells (bottom) across all treatment conditions. Both cell types exhibit normal growth patterns regardless of eVLP treatment regimen, with transient growth modulation observed between days 3-6 in T cells but complete recovery by day 9. The consistent proliferation profiles confirm that repeated eVLP exposure does not compromise long-term cellular fitness. Data represent mean ± s.e.m. from n = 2 independent experiments.

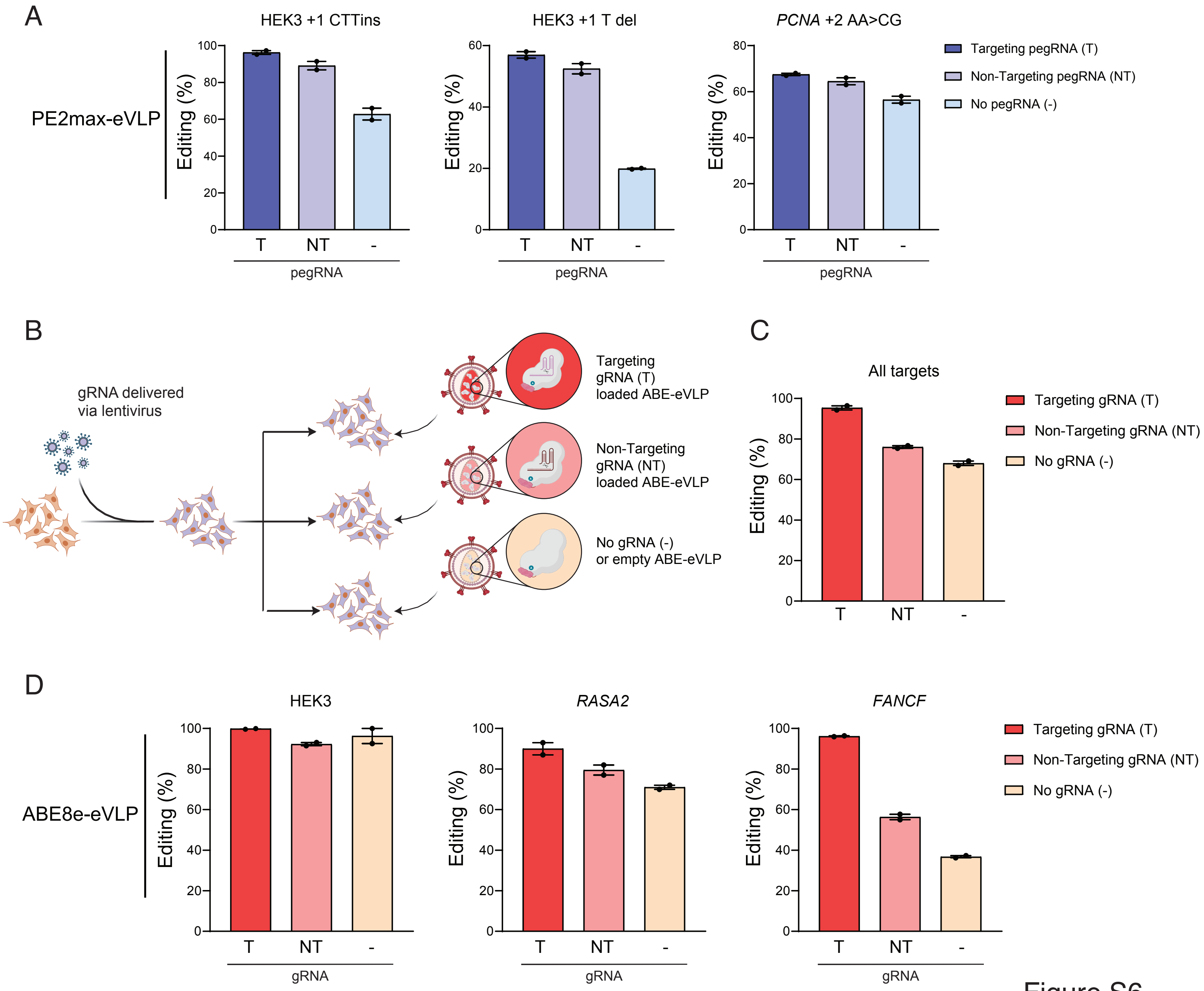

A
HEK3 +1 CTTins
HEK3 +1 T del
PCNA +2 AA>CG
PE2max-eVLP
Editing (%)
T
NT
-
pegRNA
Targeting pegRNA (T)
Non-Targeting pegRNA (NT)
No pegRNA (-)
B
gRNA delivered
via lentivirus
Targeting
gRNA (T)
loaded ABE-eVLP
Non-Targeting
gRNA (NT)
loaded ABE-eVLP
No gRNA (-)
or empty ABE-eVLP
C
All targets
Targeting gRNA (T)
Non-Targeting gRNA (NT)
No gRNA (-)
D
HEK3
RASA2
FANCF
ABE8e-eVLP
gRNA


Figure S6

**Figure S6. Validation of editor-pegRNA decoupling strategy across base editing and prime editing**

**(A)** Individual target editing efficiencies for prime editing decoupling validation. HEK293T cells transduced with lentiviral pegRNAs were subsequently treated with PE2max-eVLPs containing targeting pegRNA (T, dark blue), non-targeting pegRNA (NT, blue), or no pegRNA (-, light blue). All conditions demonstrate successful editing, confirming functional pegRNA exchange between lentiviral and eVLP delivery systems. Data represent mean ± s.e.m. from n = 2 independent experiments.

**(B)** Experimental design for base editing decoupling validation. Schematic illustrates the strategy where gRNAs are delivered via lentiviral transduction (left), while ABE editors are delivered separately using eVLPs containing three configurations: targeting gRNA (T, red), non-targeting gRNA (NT, pink), or no gRNA/empty eVLPs (-, yellow). This approach validates the generalizability of the decoupling strategy beyond prime editing.

**(C)** Summary of base editing decoupling performance averaged across all targets. Bar graph shows editing efficiency for targeting gRNA (T, 95.4%), non-targeting gRNA (NT, 76.0%), and gRNA-free (-, 68.0%) conditions, demonstrating successful editor-gRNA decoupling for adenine base editing. Data represent mean ± s.e.m. from n = 2 independent experiments.

**(D)** Individual target validation for base editing decoupling across three genomic loci (HEK3, *RASA2*, *FANCF*). This validates the broad applicability of the decoupling strategy across multiple CRISPR-based editing platforms. Data represent mean ± s.e.m. from n = 2 independent experiments.

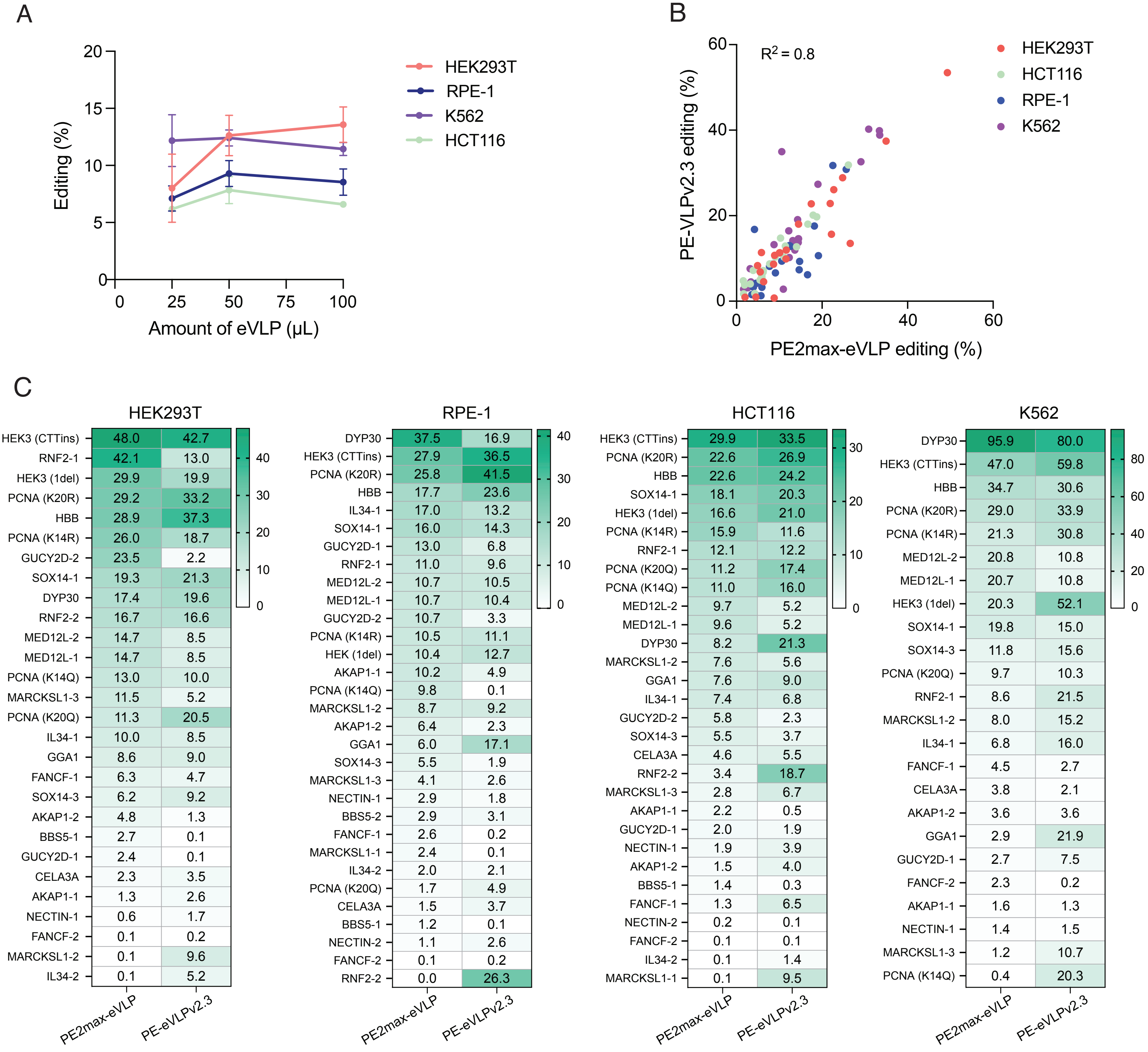
A
Editing (%)
20
15
10
5
0
0
25
50
75
100
Amount of eVLP (µL)
HEK293T
RPE-1
K562
HCT116
B
$R^2 = 0.8$
PE-VLPv2.3 editing (%)
PE2max-eVLP editing (%)
60
40
20
0
HEK293T
HCT116
RPE-1
K562
C
HEK293T
HEK3 (CTTins) 48.0 42.7
RNF2-1 42.1 13.0
HEK3 (1del) 29.9 19.9
PCNA (K20R) 29.2 33.2
HBB 28.9 37.3
PCNA (K14R) 26.0 18.7
GUCY2D-2 23.5 2.2
SOX14-1 19.3 21.3
DYP30 17.4 19.6
RNF2-2 16.7 16.6
MED12L-2 14.7 8.5
MED12L-1 14.7 8.5
PCNA (K14Q) 13.0 10.0
MARCKSL1-3 11.5 5.2
PCNA (K20Q) 11.3 20.5
IL34-1 10.0 8.5
GGA1 8.6 9.0
FANCF-1 6.3 4.7
SOX14-3 6.2 9.2
AKAP1-2 4.8 1.3
BBS5-1 2.7 0.1
GUCY2D-1 2.4 0.1
CELA3A 2.3 3.5
AKAP1-1 1.3 2.6
NECTIN-1 0.6 1.7
FANCF-2 0.1 0.2
MARCKSL1-2 0.1 9.6
IL34-2 0.1 5.2
40
30
20
10
0
PE2max-eVLP
PE-eVLPv2.3
RPE-1
DYP30 37.5 16.9
HEK3 (CTTins) 27.9 36.5
PCNA (K20R) 25.8 41.5
HBB 17.7 23.6
IL34-1 17.0 13.2
SOX14-1 16.0 14.3
GUCY2D-1 13.0 6.8
RNF2-1 11.0 9.6
MED12L-2 10.7 10.5
MED12L-1 10.7 10.4
GUCY2D-2 10.7 3.3
PCNA (K14R) 10.5 11.1
HEK (1del) 10.4 12.7
AKAP1-1 10.2 4.9
PCNA (K14Q) 9.8 0.1
MARCKSL1-2 8.7 9.2
AKAP1-2 6.4 2.3
GGA1 6.0 17.1
SOX14-3 5.5 1.9
MARCKSL1-3 4.1 2.6
NECTIN-1 2.9 1.8
BBS5-2 2.9 3.1
FANCF-1 2.6 0.2
MARCKSL1-1 2.4 0.1
IL34-2 2.0 2.1
PCNA (K20Q) 1.7 4.9
CELA3A 1.5 3.7
BBS5-1 1.2 0.1
NECTIN-2 1.1 2.6
FANCF-2 0.1 0.2
RNF2-2 0.0 26.3
40
30
20
10
0
PE2max-eVLP
PE-eVLPv2.3
HCT116
HEK3 (CTTins) 29.9 33.5
PCNA (K20R) 22.6 26.9
HBB 22.6 24.2
SOX14-1 18.1 20.3
HEK3 (1del) 16.6 21.0
PCNA (K14R) 15.9 11.6
RNF2-1 12.1 12.2
PCNA (K20Q) 11.2 17.4
PCNA (K14Q) 11.0 16.0
MED12L-2 9.7 5.2
MED12L-1 9.6 5.2
DYP30 8.2 21.3
MARCKSL1-2 7.6 5.6
GGA1 7.6 9.0
IL34-1 7.4 6.8
GUCY2D-2 5.8 2.3
SOX14-3 5.5 3.7
CELA3A 4.6 5.5
RNF2-2 3.4 18.7
MARCKSL1-3 2.8 6.7
AKAP1-1 2.2 0.5
GUCY2D-1 2.0 1.9
NECTIN-1 1.9 3.9
AKAP1-2 1.5 4.0
BBS5-1 1.4 0.3
FANCF-1 1.3 6.5
NECTIN-2 0.2 0.1
FANCF-2 0.1 0.1
IL34-2 0.1 1.4
MARCKSL1-1 0.1 9.5
30
20
10
0
PE2max-eVLP
PE-eVLPv2.3
K562
DYP30 95.9 80.0
HEK3 (CTTins) 47.0 59.8
HBB 34.7 30.6
PCNA (K20R) 29.0 33.9
PCNA (K14R) 21.3 30.8
MED12L-2 20.8 10.8
MED12L-1 20.7 10.8
HEK3 (1del) 20.3 52.1
SOX14-1 19.8 15.0
SOX14-3 11.8 15.6
PCNA (K20Q) 9.7 10.3
RNF2-1 8.6 21.5
MARCKSL1-2 8.0 15.2
IL34-1 6.8 16.0
FANCF-1 4.5 2.7
CELA3A 3.8 2.1
AKAP1-2 3.6 3.6
GGA1 2.9 21.9
GUCY2D-1 2.7 7.5
FANCF-2 2.3 0.2
AKAP1-1 1.6 1.3
NECTIN-1 1.4 1.5
MARCKSL1-3 1.2 10.7
PCNA (K14Q) 0.4 20.3
80
60
40
20
0
PE2max-eVLP
PE-eVLPv2.3

Figure S7

**Figure S7. Optimization and comparison of PE2max-eVLP systems**

**(A)** Dose-response optimization for pegRNA-free eVLP screening applications. Line graphs show editing efficiency across increasing eVLP doses (25-100 μL) in four cell lines (HEK293T, RPE-1, K562, HCT116) using the miniature sensor library. All cell lines demonstrate dose-dependent responses with saturation occurring around 50 μL, establishing the optimal dose for pooled screening applications with decoupled pegRNA delivery. Data represent mean ± s.e.m. from n = 2 independent experiments.

**(B)** Performance comparison between PE2max-eVLP (this study) and PE-eVLPv2.3 (An *et al.*, 2024[37]). Scatter plot shows strong correlation ($R^2 = 0.8$) between the two eVLP systems across four cell lines and pegRNA targets from the miniature sensor library. Each point represents one pegRNA target, with colors indicating different cell lines (HEK293T: red, HCT116: green, RPE-1: blue, K562: purple). The correlation demonstrates reproducible performance between eVLP platforms, with v2.3 showing modestly enhanced editing efficiency (1.09-fold average improvement).

**(C)** Editing efficiency heat maps across the miniature sensor library in four cell lines. Heat maps compare PE2max-eVLP (left column) versus PE-eVLPv2.3 (right column) performance for individual pegRNA targets, with color intensity representing editing efficiency (0-100% scale). Target names are listed on the y-axis, ranked by PE2max-eVLP performance. Both systems show consistent target-specific patterns across cell lines, validating the robustness of eVLP-mediated prime editing. Data represent mean values from n = 2 biological replicates per condition.

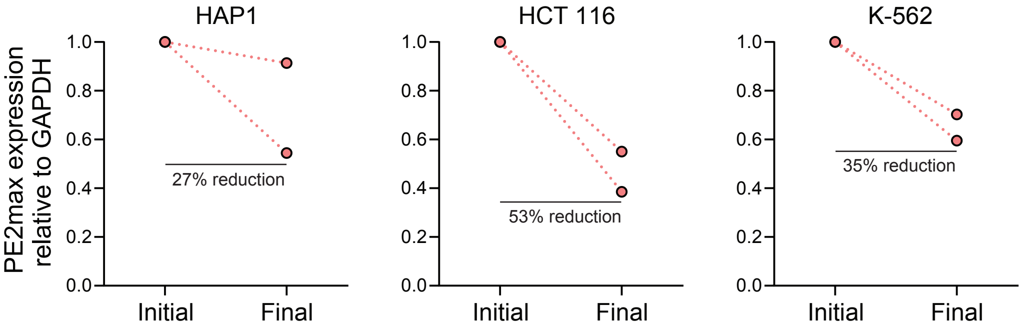
HAP1
HCT 116
K-562
PE2max expression relative to GAPDH
1.0
0.8
0.6
0.4
0.2
0.0
Initial
Final
27% reduction
53% reduction
35% reduction

Figure S8

**Figure S8. Extended analysis of PE2max silencing**

RT-qPCR analysis of PE2max transcript stability across additional cell lines. Graphs show PE2max expression levels relative to *GAPDH* housekeeping gene at initial timepoint (post-selection) and after 24 days of culture in HAP1, HCT116, and K-562 cell lines. All cell lines demonstrate significant reductions in PE2max expression over time: HAP1 (27% reduction), HCT116 (53% reduction), and K-562 (35% reduction). Individual biological replicates are shown with connecting lines, and dotted red lines indicate the trend of declining expression.

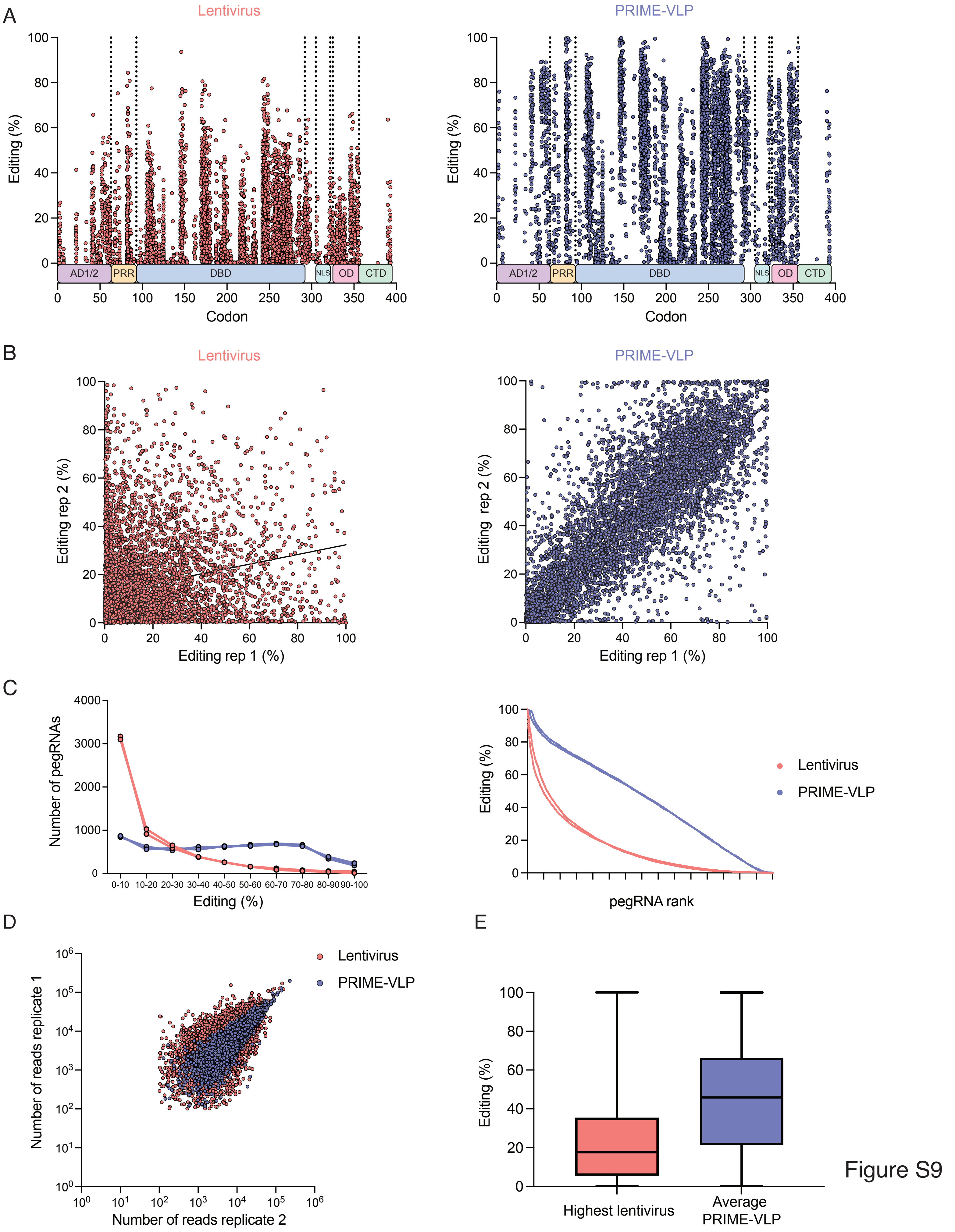
A
Lentivirus
PRIME-VLP
Editing (%)
AD1/2
PRR
DBD
NLS
OD
CTD
Codon
B
Lentivirus
PRIME-VLP
Editing rep 2 (%)
Editing rep 1 (%)
C
Number of pegRNAs
Editing (%)
pegRNA rank
Lentivirus
PRIME-VLP
D
E
Number of reads replicate 1
Number of reads replicate 2
Lentivirus
PRIME-VLP
Editing (%)
Highest lentivirus
Average PRIME-VLP

Figure S9

**Figure S9. PRIME-VLP achieves superior performance with higher reproducibility than lentiviral delivery in a high-throughput screening**

**(A)** Editing efficiency distribution across TP53 functional domains comparing lentiviral (left, red) and PRIME-VLP (right, blue) delivery. Each point represents an individual pegRNA plotted by codon position within TP53. Vertical dashed lines delineate major functional domains.

**(B)** Inter-replicate correlation analysis demonstrating superior reproducibility of PRIME-VLP. Scatter plots compare editing efficiency between biological replicates for lentiviral delivery (left, $R^2 = 0.2$) and PRIME-VLP (right, $R^2 = 0.82$). Each point represents one pegRNA. The strong correlation observed with PRIME-VLP indicates consistent editing outcomes for individual pegRNAs between replicates, while lentiviral delivery shows poor reproducibility due to stochastic expression patterns and transgene silencing effects.

**(C)** Distribution and ranking analysis of pegRNA performance. Left: Histogram showing the number of pegRNAs within defined editing efficiency ranges (0-10%, 10-20%, etc.) for lentiviral (red) and PRIME-VLP (blue) conditions. PRIME-VLP achieves a higher proportion of pegRNAs in efficient editing ranges. Right: Cumulative editing efficiency curves with pegRNAs ranked by performance, demonstrating consistently superior PRIME-VLP performance across the entire library.

**(D)** Scatter plot shows the correlation between biological replicates for sequencing read counts per pegRNA, with lentiviral delivery (red) and PRIME-VLP (blue) conditions showing comparable library representation and sequencing depth. This confirms that performance differences are not due to technical sequencing artifacts or library bias.

**(E)** Box plot compares the highest-performing lentiviral replicate for each pegRNA against the average PRIME-VLP performance. Even under these conditions that artificially favor the lentiviral approach, PRIME-VLP maintains superior editing efficiency, demonstrating the robust advantage of the eVLP-based delivery system. Box plots show median, quartiles, and range across all pegRNAs in the library.

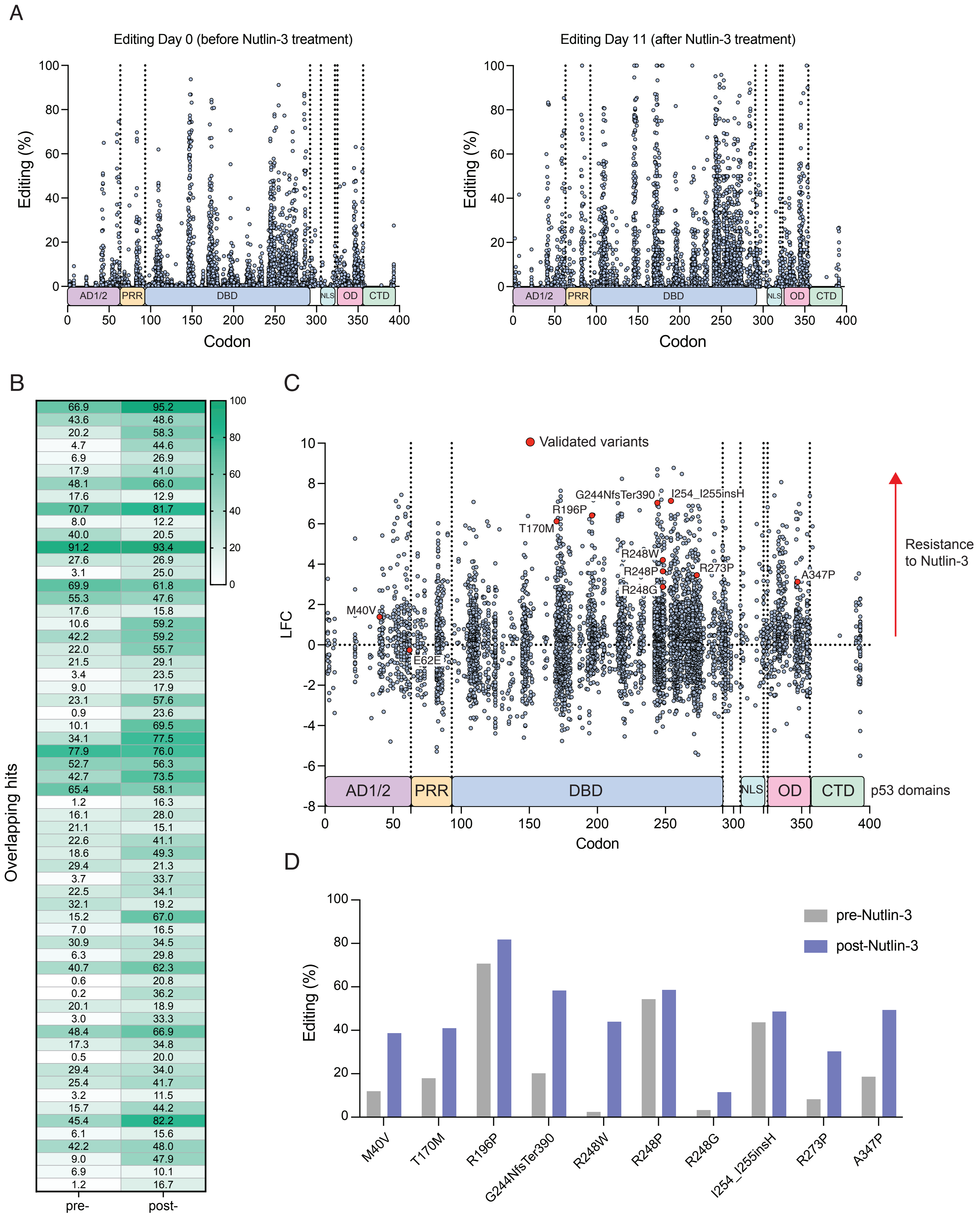
A
Editing Day 0 (before Nutlin-3 treatment)
Editing Day 11 (after Nutlin-3 treatment)
Editing (%)
Codon
AD1/2
PRR
DBD
NLS
OD
CTD
B
Overlapping hits
66.9 95.2
43.6 48.6
20.2 58.3
4.7 44.6
6.9 26.9
17.9 41.0
48.1 66.0
17.6 12.9
70.7 81.7
8.0 12.2
40.0 20.5
91.2 93.4
27.6 26.9
3.1 25.0
69.9 61.8
55.3 47.6
17.6 15.8
10.6 59.2
42.2 59.2
22.0 55.7
21.5 29.1
3.4 23.5
9.0 17.9
23.1 57.6
0.9 23.6
10.1 69.5
34.1 77.5
77.9 76.0
52.7 56.3
42.7 73.5
65.4 58.1
1.2 16.3
16.1 28.0
21.1 15.1
22.6 41.1
18.6 49.3
29.4 21.3
3.7 33.7
22.5 34.1
32.1 19.2
15.2 67.0
7.0 16.5
30.9 34.5
6.3 29.8
40.7 62.3
0.6 20.8
0.2 36.2
20.1 18.9
3.0 33.3
48.4 66.9
17.3 34.8
0.5 20.0
29.4 34.0
25.4 41.7
3.2 11.5
15.7 44.2
45.4 82.2
6.1 15.6
42.2 48.0
9.0 47.9
6.9 10.1
1.2 16.7
pre-Nutlin-3
post-Nutlin-3
Editing (%)
C
Validated variants
M40V
E62E
T170M
R196P
G244NfsTer390
I254_I255insH
R248W
R248P
R248G
R273P
A347P
Resistance to Nutlin-3
LFC
p53 domains
Codon
D
pre-Nutlin-3
post-Nutlin-3
Editing (%)
M40V
T170M
R196P
G244NfsTer390
R248W
R248P
R248G
I254_I255insH
R273P
A347P

Figure S10

**Figure S10. PRIME-VLP successfully identifies *TP53* variants and enables functional pooled prime editing screens.**

**(A)** Scatter plots compare editing efficiency distributions across *TP53* functional domains before Nutlin-3 treatment (Editing Day 0, left) and after 11 days of selection (Editing Day 11, right). Each point represents an individual pegRNA plotted by codon position within *TP53*. The overall 1.9-fold increase in editing efficiency post-selection demonstrates successful enrichment of cells harboring *TP53* loss-of-function mutations that confer resistance to p53-mediated growth arrest.

**(B)** Heat map analysis of the 62 overlaping pegRNA hits validated in previous work[21]. Each row represents a single pegRNA showing editing percentages before (Pre-Nutlin-3, left column) and after (Post-Nutlin-3, right column) Nutlin-3 treatment. Color intensity reflects editing efficiency (0-100% scale). The majority of pegRNAs show increased editing levels following selection, consistent with enrichment of functionally inactivating TP53 mutations. This demonstrates that cells harboring these variants preferentially survive Nutlin-3 treatment due to their resistance to p53-mediated cell cycle arrest.

**(C)** Comprehensive functional screen results across p53 domains. Scatter plot shows log2 fold-change (LFC) values indicating pegRNA enrichment following Nutlin-3 selection, plotted by codon position within *TP53*. Points above the dotted line (LFC = 0) represent variants conferring Nutlin-3 resistance. Previously validated variants are highlighted in red, including critical mutations such as T170M, R196P, G244NfsTer390, R248G, I254_I255insH, and A347P. The successful identification of these known functional variants confirms both the biological relevance and screening fidelity of the PRIME-VLP approach. Red arrow indicates direction of resistance to Nutlin-3.

**(D)** Individual variant analysis for selected resistance-conferring mutations. Bar graphs show editing efficiency before (grey) and after (purple) Nutlin-3 treatment for specific TP53 variants, demonstrating substantial increases in editing frequency post-selection. These include critical hotspot mutations (M40V, T170M, R196P) and known resistance variants (G244NfsTer390, R248W, R248P, R248G, I254_I255insH, R273P, A347P), confirming their functional role in conferring resistance to Nutlin-3-induced growth inhibition through p53 pathway disruption.
